# The *Burkholderia cenocepacia* type VI secretion system effector TecA is a virulence factor during lung infection

**DOI:** 10.1101/2021.07.13.451912

**Authors:** Nicole A. Loeven, Andrew I. Perault, Peggy A. Cotter, Craig A. Hodges, Joseph D. Schwartzman, Thomas H. Hampton, James B. Bliska

## Abstract

*Burkholderia cenocepacia* (Bc) is a member of the *Burkholderia cepacia* complex (Bcc), a group of bacteria with members responsible for causing lung infections in cystic fibrosis (CF) patients. The most severe outcome of Bcc infection in CF patients is cepacia syndrome, a disease characterized by necrotizing pneumonia with bacteremia and sepsis. Bc is strongly associated with cepacia syndrome making it one of the most virulent members of the Bcc. Mechanisms underlying the pathogenesis of Bc in lung infections and cepacia syndrome remain to be uncovered. Bc is primarily an intracellular pathogen, and encodes the type VI secretion system (T6SS) anti-host effector TecA, which is translocated into host cells. TecA is a deamidase that inactivates multiple Rho GTPases, including RhoA. Inactivation of RhoA by TecA triggers assembly of the pyrin inflammasome, leading to secretion of proinflammatory cytokines such as IL-1β from macrophages. Previous work with the Bc clinical isolate J2315 showed that TecA increases immunopathology during acute lung infection in C57BL/6 mice and suggested that this effector acts as a virulence factor by triggering assembly of the pyrin inflammasome. Here, we extend these results using a second Bc clinical isolate, AU1054, to demonstrate that TecA exacerbates weight loss and lethality during lung infection in C57BL/6 mice and CF mice. Unexpectedly, pyrin was dispensable for TecA virulence activity in both mouse infection models. Our findings establish that TecA is a Bc virulence factor that exacerbates lung inflammation, weight loss, and lethality in a mouse lung infection model.

**Importance:** Bc is often considered the most virulent species in the Bcc because of its close association with cepacia syndrome in addition to its capacity to cause chronic lung infections in CF patients (Loutet and Valvano 2010). Prior to this study virulence factors of Bc important for causing lethal disease had not been identified in a CF animal model of lung infection. Results of this study describe a CF mouse model and its use in demonstrating that the T6SS effector TecA of Bc exacerbates inflammatory cell recruitment and weight loss and is required for lethality and thus acts as a key virulence factor during lung infection. This model will be important in further studies to better understand TecA’s role as a virulence factor and in investigating ways to prevent or treat Bc infections in CF patients. Additionally, TecA may be the founding member of a family of virulence factors in opportunistic pathogens.

## Introduction

*Burkholderia cenocepacia* (Bc) belongs to a group of Gram-negative bacteria known as the *Burkholderia cepacia* complex (Bcc) that exist in the environment and are notorious for causing opportunistic lung infections in immunocompromised patients (Loutet and Valvano 2010; Mahenthiralingam, Urban, and Goldberg 2005). Bc infections are most commonly diagnosed in cystic fibrosis (CF) patients, however other immunocompromised individuals are also susceptible (Mahenthiralingam, Urban, and Goldberg 2005). Bcc infections are difficult to eradicate due to intrinsic antibiotic resistance (Leitão et al. 2017). Patients may present with asymptomatic, chronic, or severe infection. A severe and rapid onset form of this disease, cepacia syndrome, is characterized by necrotizing pneumonia and bacteremia, which can lead to sepsis and is often fatal in CF patients. Of the species in the Bcc, Bc accounts for ∼45% of isolates from CF patients and is most often associated with severe disease and cepacia syndrome (Hauser et al. 2011).

CF is a disease resulting from mutations in the cystic fibrosis transmembrane conductance regulator (CFTR) gene. These mutations decrease or prevent chloride ion transport, causing dehydration of the mucosal layer of the lung and gut epithelium (Ratjen et al. 2015). The most common mutation, present in at least one allele of *CFTR* in over 90% of CF patients, is the deletion of phenylalanine 508 (F508del) (Wang and Li 2014). In the CF lung dehydrated mucus prevents proper mucociliary clearance, creating a prime environment for colonization and infection predominantly by bacterial pathogens. *CFTR*, however, is not only expressed in epithelial cells. Innate immune cells also express low levels of *CFTR* and it has been shown that lack of proper chloride ion transport also affects their activities (Yoshimura et al. 1991). CF neutrophils, for example display a variety of defective responses that cause them to be hyperinflammatory yet less bactericidal (Bonfield et al. 2012; H. P. Ng, Valentine, and Wang 2016; Hang Pong Ng et al. 2014). This leads to frequent infections, predominantly bacterial, starting at a young age that begins the cycle of obstruction, infection, inflammation and exacerbation. Key bacterial pathogens in CF include *Pseudomonas aeruginosa*, *Staphylococcus aureus*, *Haemophilus influenzae*, *Stenotrophomonas maltophilia*, *Achromobacter*, and members of the Bcc (Hauser et al. 2011). Patients are often colonized with bacteria for life resulting in chronic inflammation and cumulative lung fibrosis leading to lung disease, the major cause of death of CF patients (Lee et al. 2017; Ratjen et al. 2015).

Bc is primarily but not strictly an intracellular pathogen that encodes an arsenal of virulence factors (Ganesan and Sajjan 2012; Loutet and Valvano 2010; Mahenthiralingam, Urban, and Goldberg 2005). In situ imaging of infected human and mouse lung tissue indicates that Bc resides primarily in phagocytic cells (Schwab et al. 2014; Sousa et al. 2007) but other cells may be infected including the epithelium (Sajjan et al. 2001). One factor implicated in the pathogenesis of Bc is a type VI secretion system (T6SS), designated T6SS-1 (Perault et al. 2020; Spiewak et al. 2019; Valvano 2015). T6SS-1 can deliver effectors into host cells (Valvano 2015) or other bacteria (Perault et al. 2020; Spiewak et al. 2019) that come into contact with Bc. A transposon mutagenesis screen identified T6SS-1 as an important virulence determinant in a chronic rat lung infection model (Hunt et al. 2004; Valvano 2015). Studies of macrophages infected in vitro showed that the T6SS-1 is used by intracellular Bc to inactivate the host GTPases Rac1 and Cdc42 (Valvano 2015). Inactivation of these GTPases in macrophages is associated with reduced phagocytosis and diminished assembly of NADPH oxidase complexes on phagosomes harboring Bc (Valvano 2015). Additionally, T6SS-1 was shown to be required for Bc to trigger assembly of the pyrin inflammasome in macrophages (Gavrilin et al. 2012). Canonical inflammasome assembly leads to caspase-1 processing and subsequent cleavage of gasdermin-D (GSDMD), pro-IL-1β, and pro-IL-18. The mature forms of these proinflammatory cytokines, as well as IL-1α, are subsequently released through GSDMD pores, and the infected cell may undergo pyroptosis (Kovacs and Miao 2017; Shi, Gao, and Shao 2017; Xu et al. 2014). Xu et al (Xu et al. 2014) discovered that inactivation of Rho GTPases (RhoA/B/C) in macrophages infected with Bc in a T6SS-1-dependent manner is responsible for activation of the pyrin inflammasome. This work established that pyrin, encoded by the *Mefv* gene, is an intracellular inflammasome sensor, that specifically and indirectly senses inactivation of Rho A/B/C (hereafter referred to as RhoA) by bacterial effectors and toxins (Loeven, Medici, and Bliska 2020; Xu et al. 2014; De Zoete and Flavell 2014). Pyrin is the only inflammasome sensor that it is preferentially expressed in phagocytes such as neutrophils, monocytes and activated macrophages (Schnappauf et al. 2019).

TecA (<u>T</u>6SS <u>e</u>ffector protein affecting <u>c</u>ytoskeletal <u>a</u>rchitecture) was subsequently identified as an enzyme encoded by Bc that is injected into host phagocytes and inactivates RhoA and Rac 1 (Aubert et al. 2016). Additionally, Aubert et al. determined that TecA deamidates asparagine-41 of RhoA, triggering pyrin activation and inflammasome assembly (Aubert et al. 2016). Thus, pyrin inflammasome assembly in response to inactivation of RhoA by TecA is an effector-triggered immune response that is mechanistically similar to the “guard hypothesis” in plant immunity (Aubert et al. 2016). This work also identified *tecA* genes in other Bc strains and *tecA*-like gene sequences in other bacteria (Aubert et al. 2016). A putative protein structure for TecA was generated which led to a prediction that cysteine-41 comprises part of a catalytic triad (Aubert et al. 2016). A codon change mutation of cysteine-41 to alanine (C41A) in *tecA* resulted in loss of catalytic function for TecA and prevented pyrin inflammasome activation in Bc-infected macrophages (Aubert et al. 2016).

Studies using acute lung infection models in mice have been carried out with the Bc strain J2315 (BcJ2315) to understand the role of pyrin and TecA in pathogenesis. Xu et al. found by histopathology that an increase in inflammation seen in C57BL/6 mice infected with BcJ2315 was not observed in isogenic *Mefv^-/-^* mice, suggesting that early immune cell influx to the lungs, and subsequent immunopathology, was pyrin-dependent (Xu et al. 2014). Results of a similar infection procedure showed that TecA catalytic activity was required for BcJ2315 to cause immune cell recruitment and injury to the lungs of C57BL/6 mice (Aubert et al. 2016). Although these results suggest that TecA acts as a virulence factor by triggering assembly of the pyrin inflammasome, leading to inflammatory cell recruitment and lung immunopathology, other measurements of disease (e.g. pathogen burdens, lethality) have not been examined to confirm this concept. Additionally, the roles of pyrin and TecA in Bc pathogenesis have only been investigated with BcJ2315 and have not been studied in mice lacking Cftr function, which are known have increased susceptibility to lung infections (Abdulrahman et al. 2011; Robledo-avila et al. 2018; Sajjan et al. 2001). Here we used Bc strain AU1054 (BcAU1054), isolated from the bloodstream of a CF patient in the US (Perault et al. 2020), to perform the first study on pyrin and TecA and their virulence roles during lung infection of C57BL/6 mice and mice with the F508del mutation in *Cftr*.

## Results

### TecA is a virulence factor during BcAU1054 lung infection in WT mice

To study Bc pathogenesis and immunity we established a mouse lung infection model with BcAU1054 (Table 1), a genomovar IIIB, PHDC lineage clinical isolate (Perault et al. 2020). In initial experiments C57BL/6 (WT) mice were mock infected or infected with 5×10^7^ CFU of BcAU1054 by oropharyngeal (o.p.) aspiration and at 12 hours post infection immunohistochemistry (IHC) was performed on lung sections using mouse antisera to Bc. As shown in Fig. S1, Bc antisera staining was only detected in infected sections and was concentrated in what appeared to be phagocytes primarily located in the alveoli. This IHC staining pattern for Bc in phagocytes is similar to what has been reported for infected human and mouse lung sections (Schwab et al. 2014; Sousa et al. 2007).

**Table 1.**
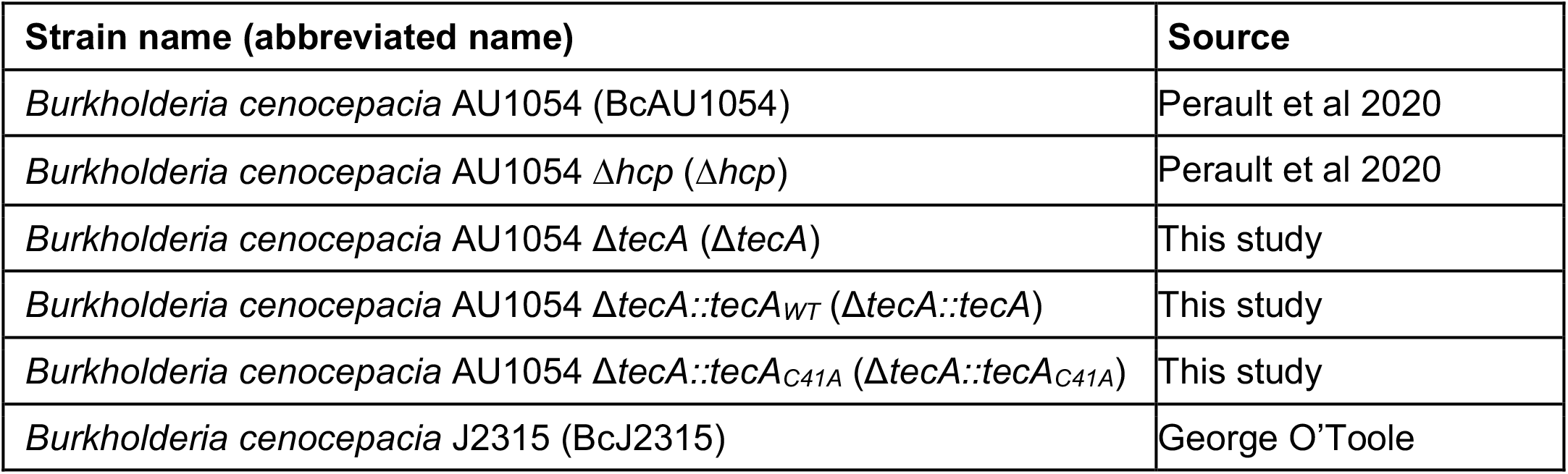
Bacterial strains used in this study

| Strain name (abbreviated name) | Source |
| --- | --- |
| *Burkholderia cenocepacia* AU1054 (BcAU1054) | Perault et al 2020 |
| *Burkholderia cenocepacia* AU1054 $\Delta hcp$ ($\Delta hcp$) | Perault et al 2020 |
| *Burkholderia cenocepacia* AU1054 $\Delta tecA$ ($\Delta tecA$) | This study |
| *Burkholderia cenocepacia* AU1054 $\Delta tecA::tecA_{WT}$ ($\Delta tecA::tecA$) | This study |
| *Burkholderia cenocepacia* AU1054 $\Delta tecA::tecA_{C41A}$ ($\Delta tecA::tecA_{C41A}$) | This study |
| *Burkholderia cenocepacia* J2315 (BcJ2315) | George O'Toole |

To study the role of TecA in pathogenesis, *tecA* was deleted from BcAU1054 (Δ*tecA*) (Table 1). Groups of WT mice were then infected with BcAU1054 or Δ*tecA* and weight and survival was monitored for 14 days. Compared to mice infected with Δ*tecA*, Bc AU1054-infected mice experienced significant weight loss (Fig. 1A). The weight loss data in Fig. 1A was used to estimate the average change in weight from the previous day (log2) and results were displayed by locally weighted scatterplot smoothing (LOESS) (data not shown). Using the Asymptotic Wilcoxon-Mann-Whitney Test, it was determined that the mice infected with Δ*tecA* regained weight significantly earlier as compared to BcAU1054 (p-value = 0.007201). Fifty percent of the mice infected with BcAU1054 succumbed to the disease while Δ*tecA* was significantly less virulent (Fig. 1B), indicating that TecA is a virulence factor. In different sets of experiments the percentages of WT mice that died from BcAU1054 infection ranged from 50% (Fig. 1B) to 20% (see Fig. S4B) while Δ*tecA* was reproducibly avirulent. To demonstrate that the avirulent phenotype of Δ*tecA* was due to loss of TecA enzymatic activity, we complemented Δ*tecA* with wild-type *tecA* (Δ*tecA::tecA_WT_*) or catalytically-inactive *tecA* (Δ*tecA::tecA_C41A_*) (Table 1) and carried out mouse infections with the resulting strains. The avirulent phenotype of Δ*tecA* was complemented by *tecA*_WT_ but not *tecA_C41A_* as determined by the weight loss assay (Fig. S2) demonstrating that the enzymatic activity of TecA is required for virulence. To examine how TecA promotes virulence, CFU assays were performed on lungs and spleens of mice infected with BcAU1054 or Δ*tecA* over a time course. There was no trend indicating a difference in the numbers of BcAU1054 and Δ*tecA* CFUs detected over the time course up to day 4 (Fig. 1C,D), suggesting that TecA does not increase the burden of Bc in the lung or dissemination to the spleen.

**Fig. 1.**
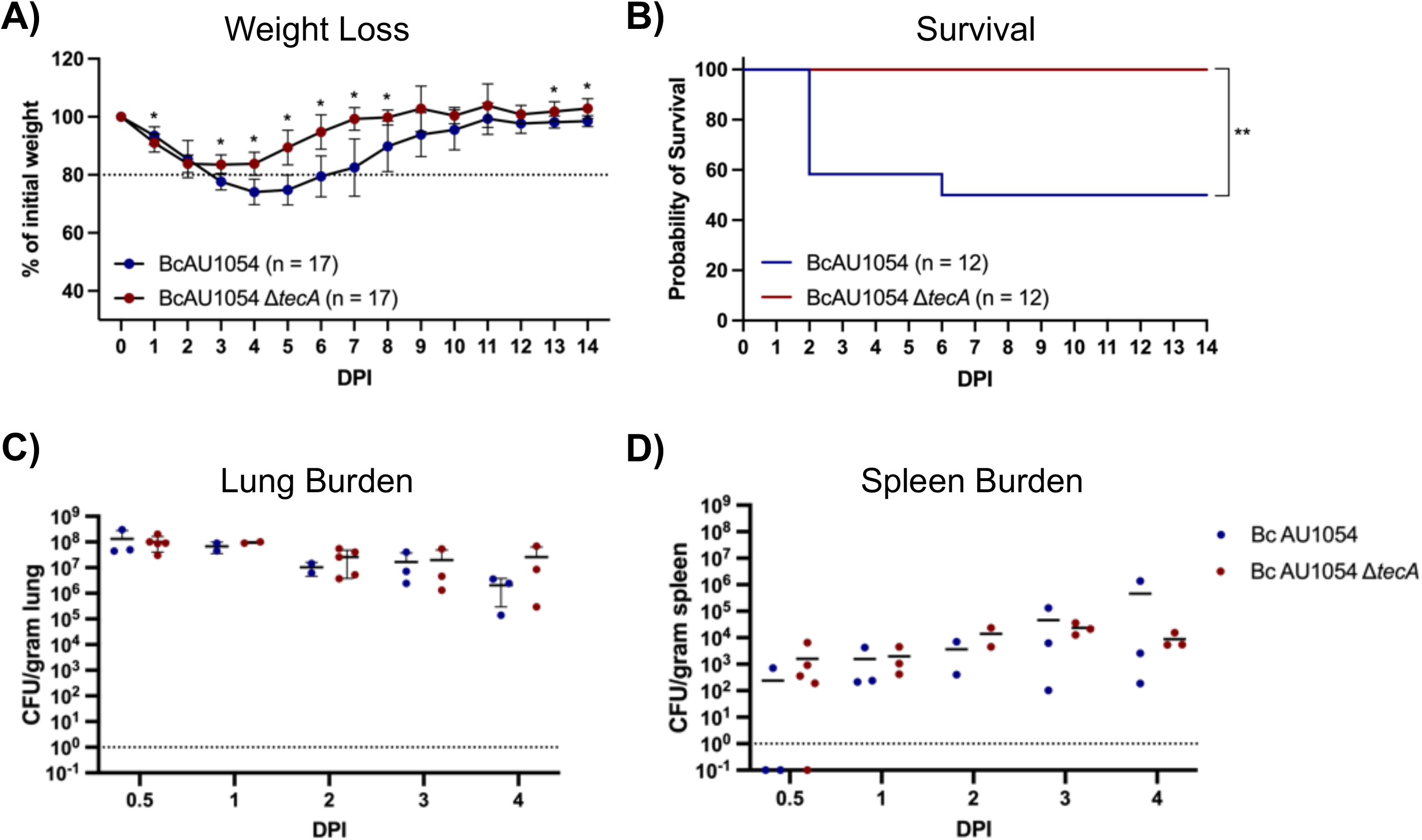
TecA enhances weight loss and lethality in WT mice infected with BcAU1054. A) Weight loss of WT mice infected by o.p. instillation with 5×10^7^ CFU of BcAU1054 or Δ*tecA*. Mice were weighed on D_0_ and daily for 14 days post infection (DPI). Results were pooled from 3 independent experiments, n = total mice infected. Data are represented as percent of initial weight. Days on which the two groups differed significantly (p < .05, Welch’s T test) are marked with an asterisk. B) Percent survival monitored for 14 DPI of WT mice infected with BcAU1054 or Δ*tecA*. Results were pooled from 3 independent experiments, n = total mice infected. **, p = 0.0054, by Log-Rank (Mantel-Cox) test. C) Lung and D) spleen burdens at the indicated DPI in WT mice infected with BcAU1054 or Δ*tecA*. Results pooled from multiple experiments, each point represents a value obtained from an individual mouse and data are represented as CFU per gram of tissue.

Staining of lung sections with hematoxylin and eosin (H&E) showed mild edema and peri-bronchiolar and alveolar inflammatory infiltrates that appeared more extensive in the BcAU1054 infections as compared to Δ*tecA*, which had regions that appeared essentially uninvolved, at both 12 hours (Fig. 2A,B) and 3 days (Fig. 2C,D) post challenge. These results are in line with those of Aubert et al. who used a similar infection model with BcJ2315 in WT mice and showed that TecA increases infiltration of inflammatory cells and lung damage at 12 hours (Aubert et al. 2016). Semi-quantitative scoring of blinding images by a pathologist indicated that infection by both BcAU1054 and Δ*tecA* caused similar neutrophilic and mononuclear cell infiltration and damage in the bronchioles and alveoli (Table S1). To better quantify the cell infiltration IHC analysis of lung sections using antibodies to CD11b, Ly6G and F4/80 was carried out. Representative images are shown in Fig. 3A and B, and quantitative analysis results normalized using H-scores are shown in Fig. 3C-E. Results showed that BcAU1054 TecA enhanced neutrophil (CD11b^+^ Ly6G^+^) infiltration at 12 hours post infection (Fig. 3A,C,D) and inflammatory monocyte-derived macrophage (CD11b^+^ F4/80^+^) infiltration at 3 days post infection (Fig. 3B,C,E). By 3 days post infection the neutrophil numbers in the lungs had decreased to background levels in both infection conditions (Fig. 3B,D) while numbers of inflammatory monocyte-derived macrophages had increased at this time point in response to infection with BcAU1054 (Fig. 3B,C,E). These results are consistent with BcAU1054 TecA exacerbating an acute lethal bronchopneumonia that is dominated by a sustained increase in F4/80^+^ inflammatory monocyte-derived macrophages at day 3 post infection.

**Fig. 2.**
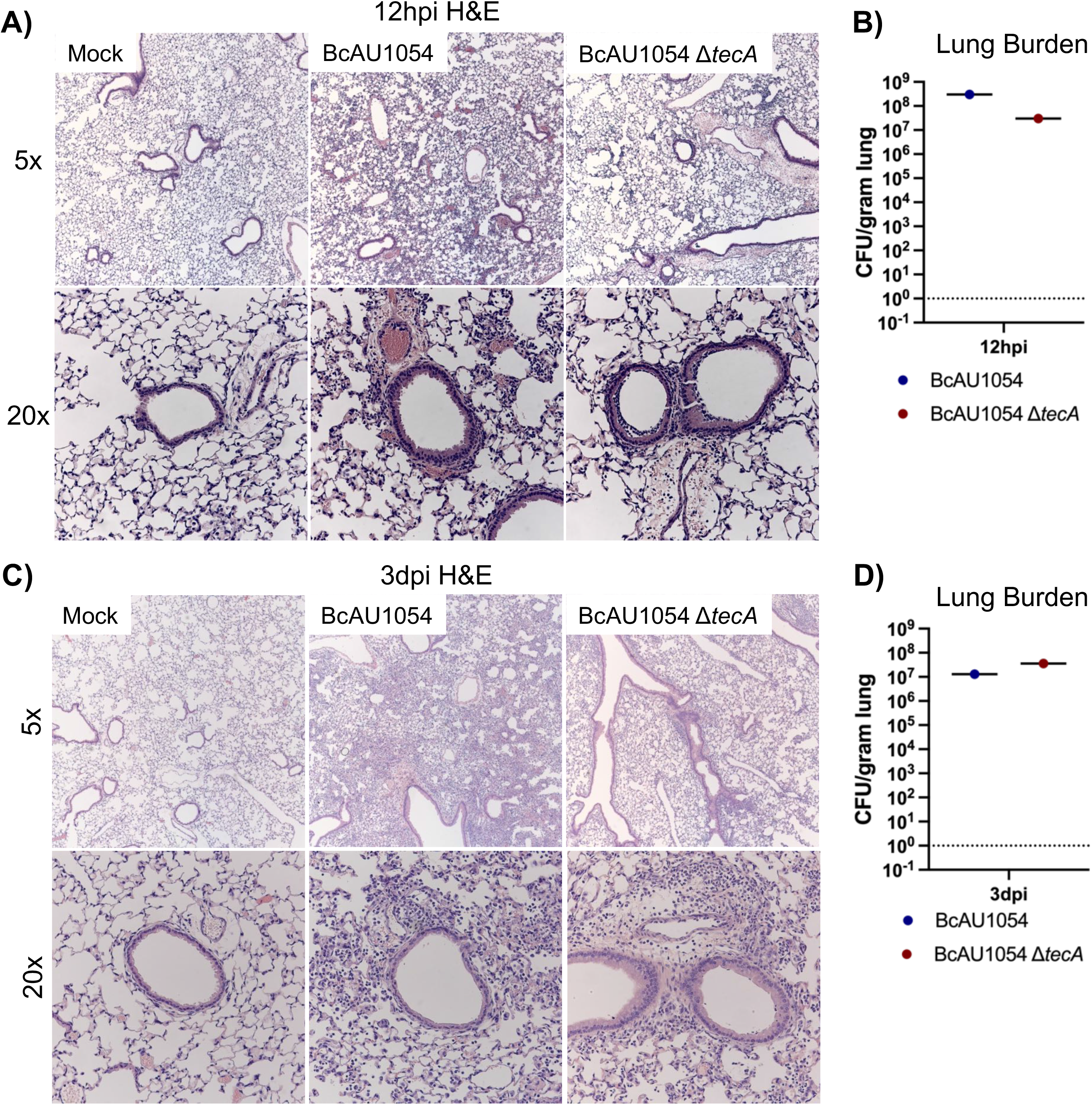
Lung inflammation in WT mice infected with BcAU1054 or Δ*tecA*. A) Representative H&E images of the four right lung lobe sections from a WT mouse left uninfected (mock) or infected o.p. with 5×10^7^ CFU of BcAU1054 or Δ*tecA* at A) 12 hpi and C) 3 dpi. Light microscopy images are shown at 5X and 20X magnification. Burdens of BcAU1054 or Δ*tecA* in the infected left lungs of the mice analyzed in A) and C) are shown in B) and D) respectively. Data are represented as CFU per gram of lung.

**Fig. 3.**
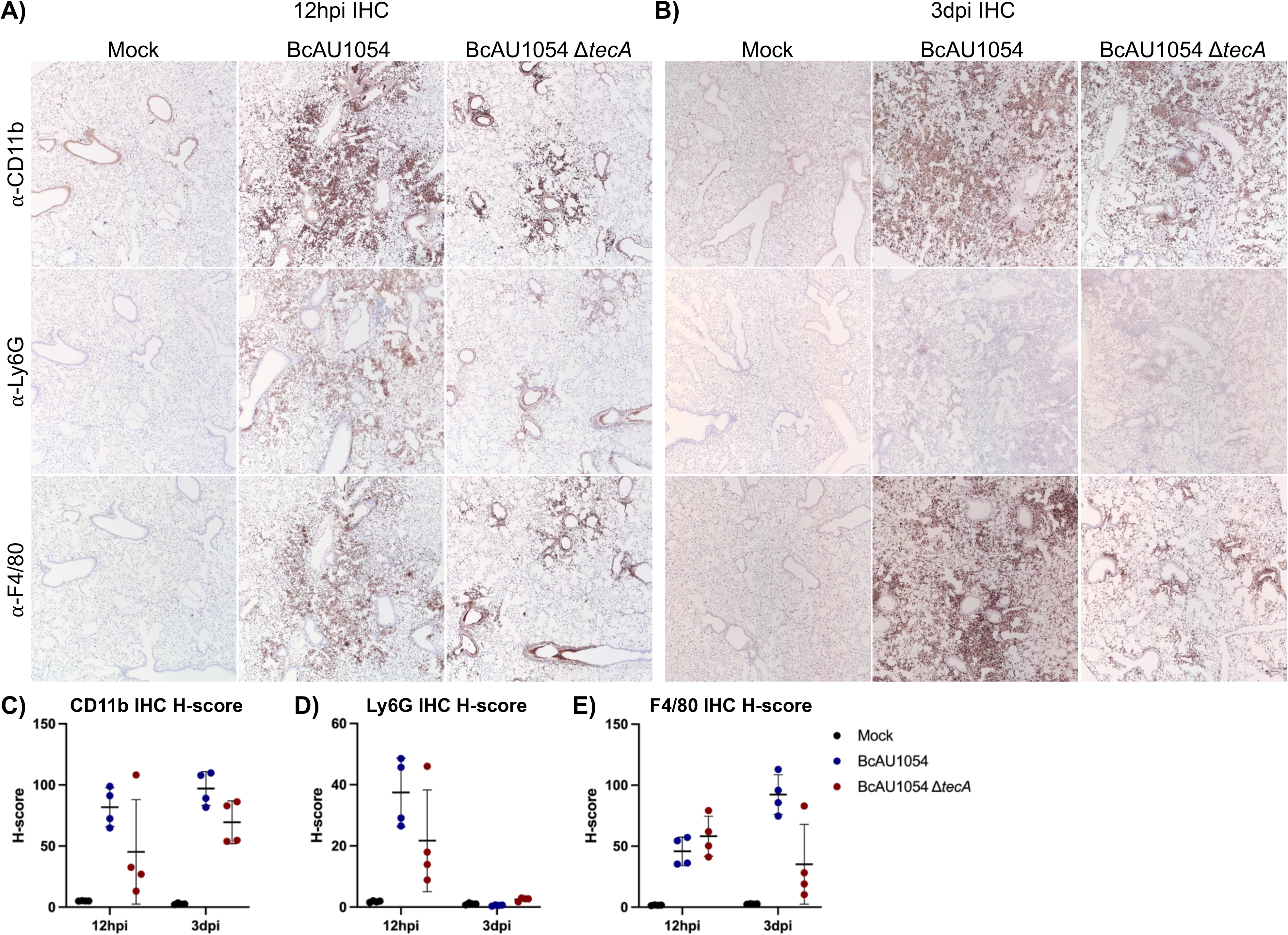
TecA enhances recruitment of neutrophils and inflammatory monocyte-derived macrophages to lungs in WT mice infected with BcAU1054. IHC was performed on sections from the right lung lobes analyzed in Fig. 2 using CD11b, Ly6G and F4/80 antibodies as indicated. Representative IHC images captured by light microscopy at 5X magnification are shown. Quantitative IHC image analysis results normalized using H-scores are shown in C) for CD11b, D) for Ly6G and E) for F4/80. Each data point represents the H-score for each of the four different lobe sections.

### TecA is a virulence factor during BcAU1054 lung infection in CF mice

Previous studies have indicated that mice lacking Cftr function have increased susceptibility to Bc lung infections (Abdulrahman et al. 2011; Robledo-avila et al. 2018; Sajjan et al. 2001). To examine this possibility in our infection model with BcAU1054, *Cftr^F508del^* (*Cftr^em1Cwr^*) mice were obtained from the Case Western Reserve University CF Mouse Model Core. *Cftr^F508del^* mice were infected with BcAU1054 or Δ*tecA* as above. *Cftr^F508del^* mice infected with BcAU1054 had increased weight loss as compared to Δ*tecA* (Fig. 4A) although this difference was not significant when the data was displayed by LOESS and analyzed using the Asymptotic Wilcoxon-Mann-Whitney Test (data not shown). TecA was required for lethality (Fig. 4B), without impacting lung or spleen CFU over the first 3 days of infection (Fig. 4C,D). In different sets of experiments the percentages of *Cftr^F508del^* mice that died from BcAU1054 infection ranged from 80% (Fig. 4B) to ∼50% (see Fig. 7B), suggesting that the *Cftr^F508del^* mice are more susceptible to lethal disease as compared to WT mice. BcAU1054 and Δ*tecA* infection increased inflammation in the lungs of *Cftr^F508del^* mice at 12 hours and 3 days post infection, and the H&E staining results were similar to those obtained in WT mice (Fig. 5, compare with Fig. 2). Lung sections from 12 hours post infection and 3 days post infection were also stained for IHC using antibodies to CD11b, Ly6G or F4/80 (Fig. 6). Overall, the staining pattern was similar to that in WT mice, although there was no trend toward increased Ly6G^+^ neutrophils in the BcAU1054- as compared to Δ*tecA*-infected lung sections at 12 hours (Fig. 6D, compare with Fig. 3D). In summary, BcAU1054 TecA functioned as a virulence factor during lung infection in *Cftr^F508del^* mice.

**Fig. 4.**
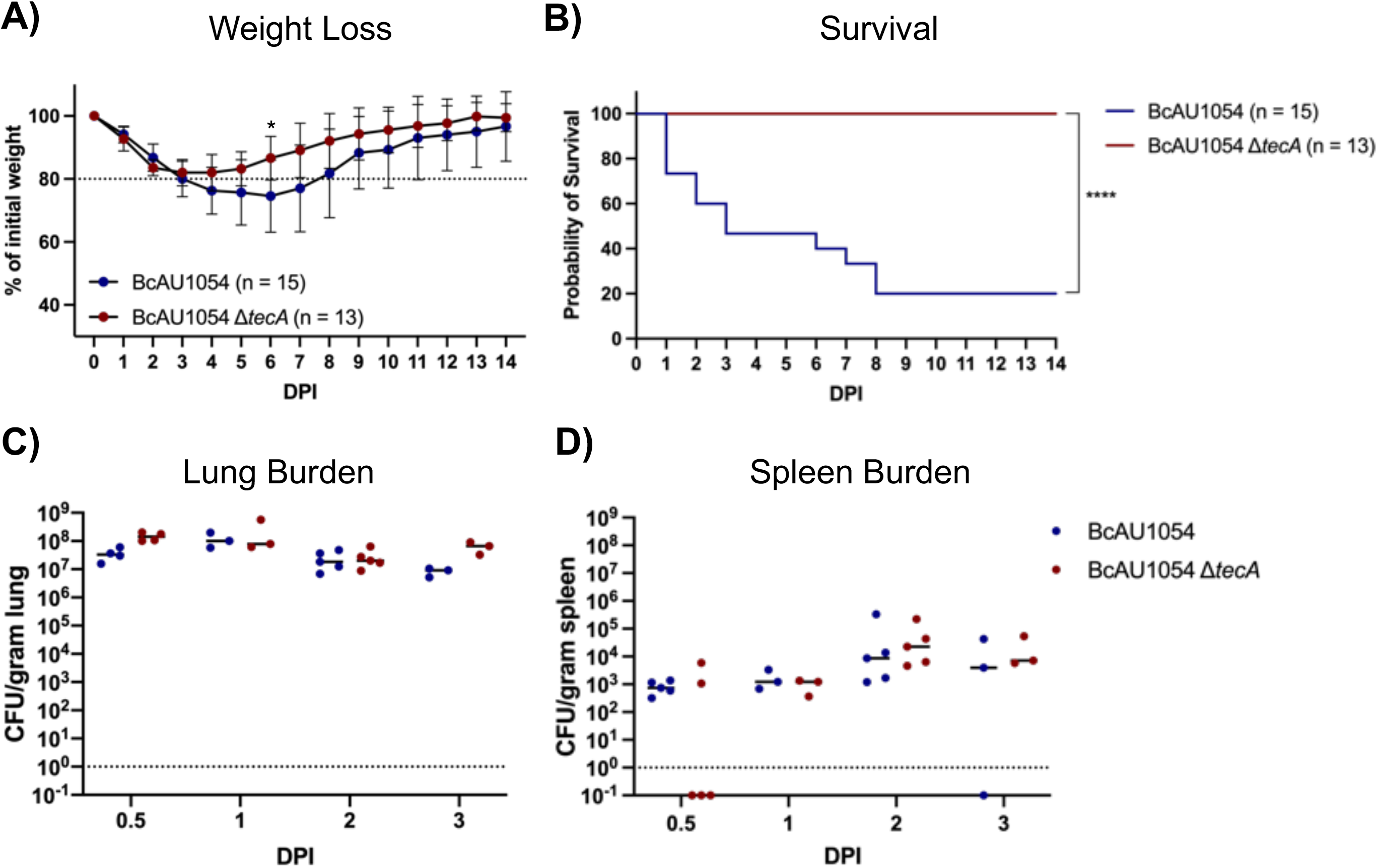
TecA enhances weight loss and lethality in *Cftr^F508del^* mice infected with BcAU1054. A) Weight loss of *Cftr^F508del^* mice infected o.p. with 5×10^7^ CFU of BcAU1054 or Δ*tecA*. Mice were weighed on D_0_ and daily for 14 DPI. Results were pooled from 5 independent experiments, n = total mice infected. Data are represented as percent of initial weight. Days on which the two groups differed significantly (p < .05, Welch’s T test) are marked with an asterisk. B) Percent survival monitored for 14 DPI of *Cftr^F508del^* mice infected with BcAU1054 or Δ*tecA*. Results were pooled from five independent experiments, n = total mice infected. ****, p = <0.0001, by Log-Rank (Mantel-Cox) test. C) Lung and D) spleen burden at the indicated DPI in *Cftr^F508del^* mice infected with BcAU1054 or Δ*tecA*. Results pooled from multiple experiments, each point represents a value obtained from an individual mouse. Data are represented as CFU per gram of tissue.

**Fig. 5.**
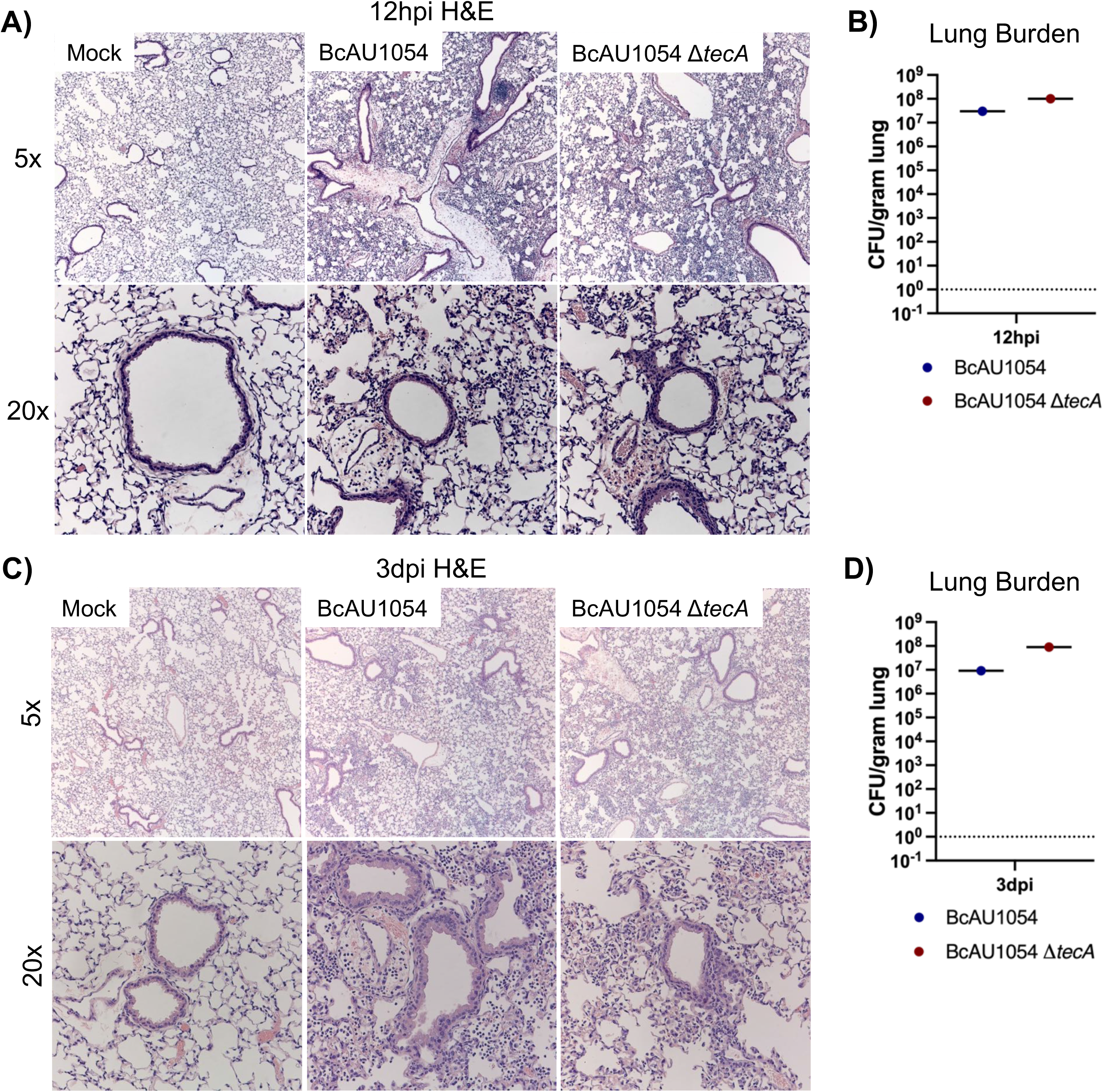
Lung inflammation in *Cftr^F508del^* mice infected with BcAU1054 or Δ*tecA*. A) Representative H&E images of the four right lung lobe sections from a *Cftr^F508del^* mouse left uninfected (mock) or infected o.p. with 5×10^7^ CFU of BcAU1054 or Δ*tecA* at A) 12 hpi or C) 3 dpi. Light microscopy images are shown at 5X and 20X magnification. Burdens of BcAU1054 or Δ*tecA* in the infected left lungs of the mice analyzed in A) and C) are shown in B) and D) respectively. Data are represented as CFU per gram of lung.

**Fig. 6.**
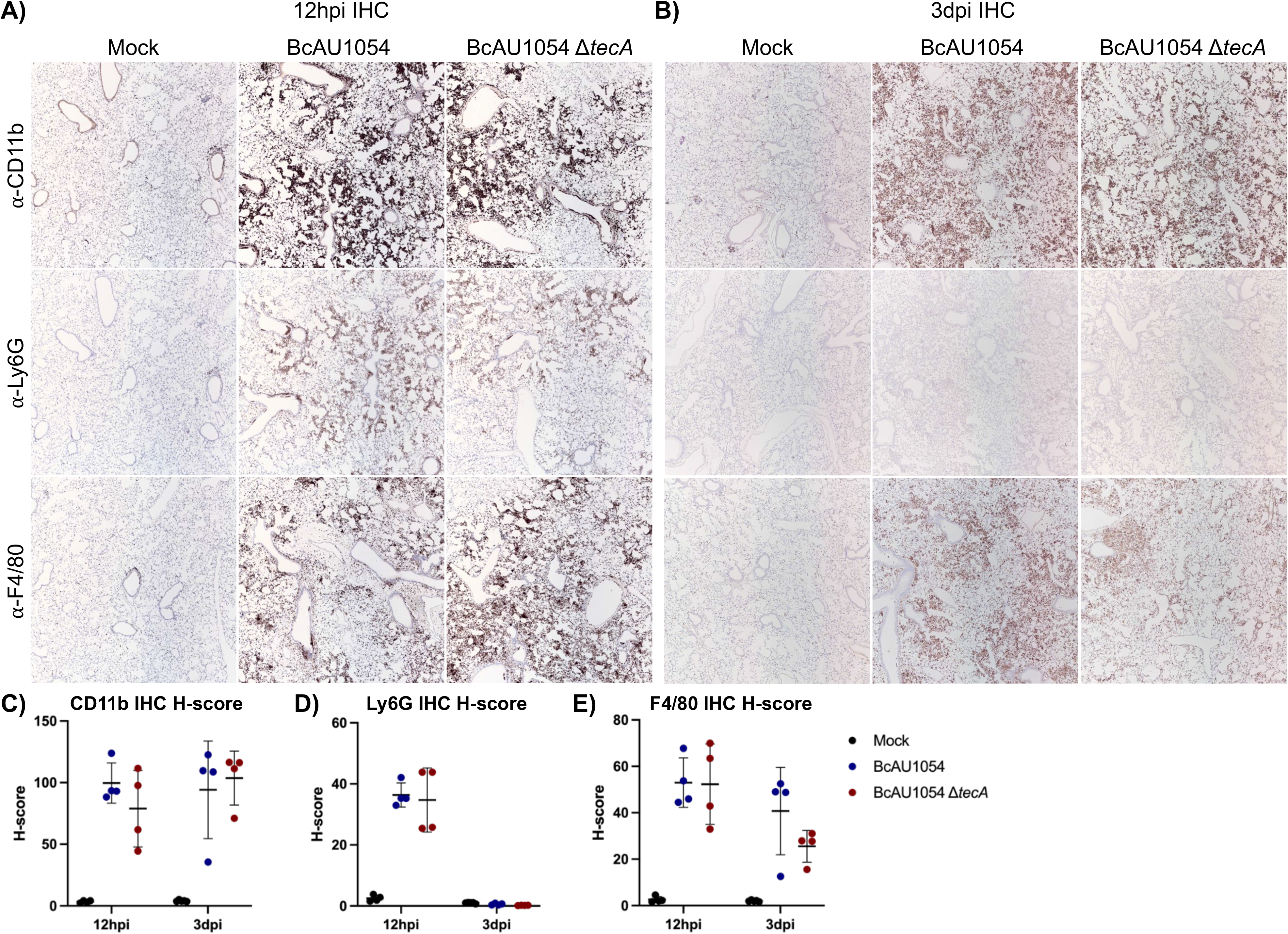
TecA enhances recruitment of inflammatory monocyte-derived macrophages to lungs in *Cftr^F508del^* mice infected with BcAU1054. IHC was performed on sections from the right lung lobes analyzed in Fig. 5 using CD11b, Ly6G and F4/80 antibodies as indicated. Representative IHC images captured by light microscopy at 5X magnification are shown. Quantitative IHC image analysis results normalized using H-scores are shown in C) for CD11b, D) for Ly6G and E) for F4/80. Each data point represents the H-score for each of the four different lobe sections.

**Fig. 7.**
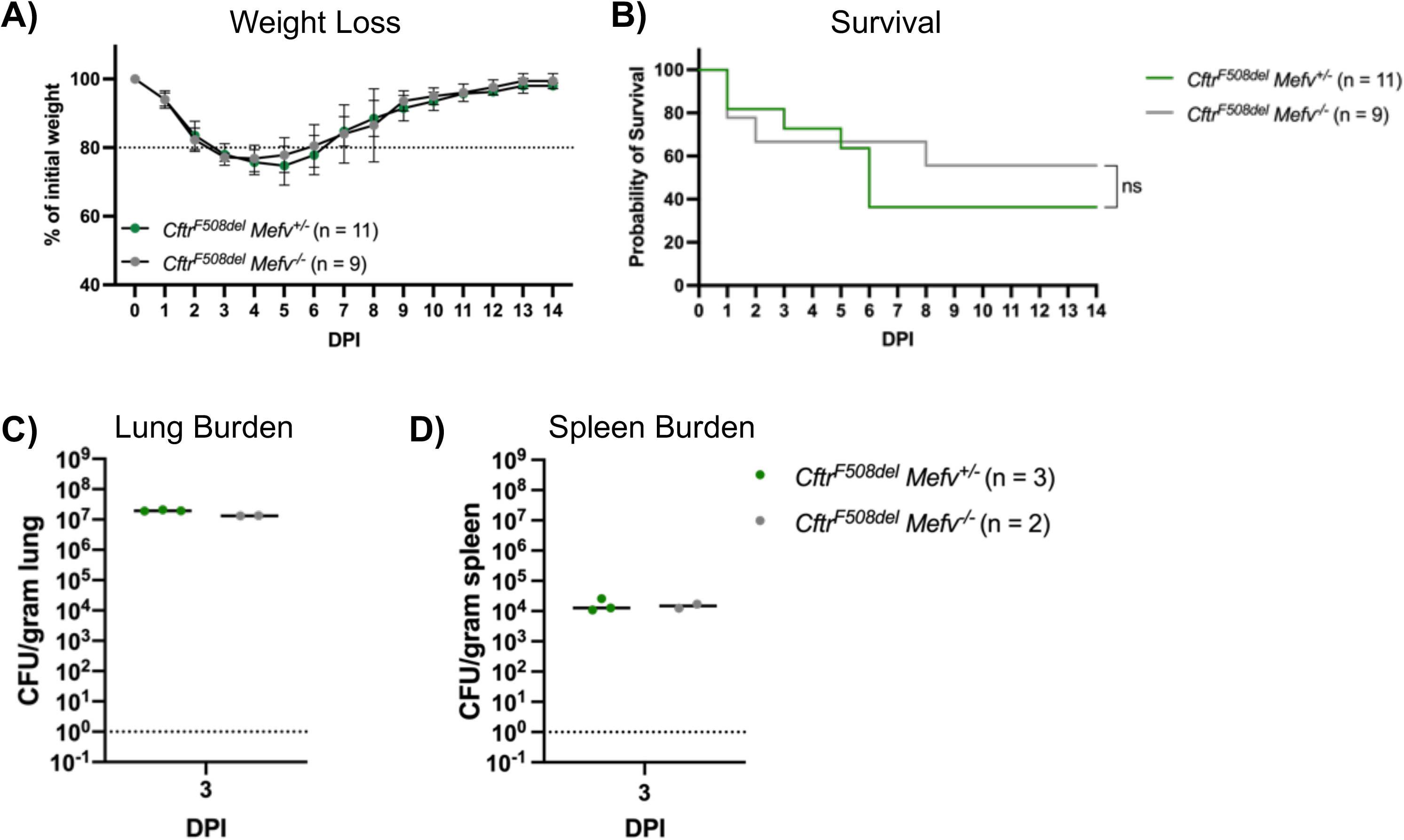
Pyrin is dispensable for BcAU1054 virulence in *Cftr^F508del^* mice. A) Weight loss of *Cftr^F508del^ Mefv^+/-^* and *Cftr^F508del^ Mefv^-/-^* mice infected o.p. with 5×10^7^ CFU of BcAU1054. Mice were weighed on D_0_ and daily for 14 DPI. Results were pooled from 3 independent experiments, n = total mice infected. Data are represented as percent of initial weight. B) Percent survival monitored for 14 days post infection (DPI) of *Cftr^F508del^ Mefv^+/-^* and *Cftr^F508del^ Mefv^-/-^* mice infected with BcAU1054. Results were pooled from three independent experiments, n = total mice infected. ns, not significant by Log-Rank (Mantel-Cox) test. C) Lung and D) spleen burden at 3 DPI in *Cftr^F508del^ Mefv^+/-^* and *Cftr^F508del^ Mefv^-/-^* mice infected with BcAU1054. Data are represented as CFU per gram of tissue.

### Pyrin is dispensable for Bc virulence during lung infection in mice

Having established that TecA is a virulence factor, experiments were carried out to determine the importance of the pyrin inflammasome for pathogenesis during BcAU1054 lung infection in mice. To confirm that TecA in BcAU1054 can trigger pyrin inflammasome assembly in WT and *Cftr^F508del^* backgrounds, *in vitro* infection experiments were carried out using LPS-primed bone marrow derived macrophages (BMDMs). WT and *Cftr^F508del^* BMDMs were left uninfected or infected at an MOI of 20 for 90 minutes with BcAU1054, Δ*tecA*, Δ*tecA::tecA_WT_* or Δ*tecA::tecA_C41A_*. Purified TcdB from *Clostridium difficile* was used as a positive control for pyrin inflammasome activation. Results of western blotting and ELISA were used to monitor the outcomes of the infections or intoxications. In its inactive conformation murine pyrin is phosphorylated at serine 205 (p-S205) and serine 241 (Park et al. 2016). Dephosphorylation of these sites appears to trigger activation of pyrin (Gao et al. 2016; Park et al. 2016). The phosphorylation status of pyrin was determined by western blotting for total pyrin and p-S205 pyrin (Gao et al. 2016). WT and *Cftr^F508del^* BMDMs infected with strains BcAU1054 or Δ*tecA::tecA_WT_* or intoxicated with TcdB had decreased levels of p-S205 pyrin whereas those infected with Δ*tecA,* or Δ*tecA::tecA_C41A_* retained p-S205 (Fig. S3A). Pro-IL-1β was detected by western blotting at similar levels in LPS-primed BMDMs in all conditions (Fig. S3A). ELISA results showed that the pyrin inflammasome was assembled in response to the activities of TecA or TcdB similarly in WT or *Cftr^F508del^* BMDMs, resulting in equivalent processing and release of IL-1β by both cell types (Fig. S3B).

The *Cftr^F508del^* mice were crossed to *Mefv^-/-^* mice and the offspring were infected with BcAU1054 to investigate the role of the pyrin inflammasome for pathogenesis during BcAU1054 lung infection. As shown in Fig. 7, there was no difference in weight loss (A), survival (B) or lung (C) or spleen (D) CFU burdens (day 3), respectively, when *Cftr^F508del^ Mefv^+/-^* or *Cftr^F508del^ Mefv^-/-^* mice were infected with BcAU1054. To determine if *Mefv^+/-^* mice produce pyrin and secrete IL-1β to a similar extent as the *Mefv^+/+^* background, *Cftr^F508del^ Mefv^+/+^*, *Cftr^F508del^ Mefv^+/-^* and *Cftr^F508del^ Mefv^-/-^* BMDMs were infected as described above with BcAU1054, Δ*tecA,* or the Δ*hcp* T6SS-1-deficient mutant (Table 1) or intoxicated with TcdB. Results showed that uninfected *Cftr^F508del^ Mefv^+/-^* BMDMs produce pyrin at the expected ∼2-fold lower amounts compared to *Cftr^F508del^ Mefv^+/+^* BMDMs (Fig. S3C) yet IL-1βrelease triggered by TecA or TcdB was similar (Fig. S3D).

To extend these results to the non-CF background, WT or *Mefv^-/-^* mice were infected with BcAU1054. As shown in Fig. S4, there was no difference in weight loss (A), survival (B) or organ CFU burdens at day 3 (C,D) when WT or *Mefv^-/-^* mice were infected with BcAU1054. Lung sections from these mice infected with BcAU1054 were examined after H&E staining and no difference in inflammatory cell infiltration was evident between WT and *Mefv^-/-^* genotypes (Fig. S5). Production of pyrin in inflammatory cells obtained by bronchoalveolar lavage (BAL) of WT mice infected with BcAU1054 or BcAU1054Δ*tecA* was confirmed by immunoblotting (Fig. S6). Our results with BcAU1054 (genomovar IIIB, PHDC lineage) thus differed from Xu et al., who reported that inflammatory cell infiltration into lungs of mice intranasally infected with BcJ2315 (genomovar IIIA, ET12 lineage) required the pyrin inflammasome (Xu et al. 2014).

To determine if the discrepancy between our results and those of Xu et al. was due to Bc strain differences, *Mefv^+/-^* and *Mefv^-/-^* mice were infected o.p. with 5×10^7^ CFU of BcJ2315 (Table 1) (Xu et al. 2014). BcJ2315 appeared to be less virulent than BcAU1054 as the weight loss was less severe in the infected mice (Fig. S7A). Nevertheless, there was no difference in weight loss or survival for *Mefv^+/-^* and *Mefv^-/-^* mice infected with BcJ2315 (Fig. S7A,B). In addition, H&E-stained lung sections from *Mefv^+/-^* and *Mefv^-/-^* showed no difference in inflammatory cell infiltration at 12 hours and 3 days post infection with BcJ2315 (Fig. S8A, C). We confirmed that our isolate of BcJ2315 triggered assembly of the pyrin inflammasome in LPS-primed BMDMs infected in vitro (Fig. S3E). Thus, the requirement for pyrin in lung inflammation previously reported with the BcJ2315 strain (Xu et al. 2014) was not reproduced in our mouse infection model.

## Discussion

The underlying basis for the pathogenesis of Bc during lung infection remains to be fully understood. Aubert et al. demonstrated that TecA is critical for BcJ2315 to cause inflammatory cell recruitment and damage to lungs in WT mice (Aubert et al. 2016). We have extended these results in the following ways: 1) We show that TecA is virulence factor that exacerbates weight loss and lethality in addition to inflammatory cell recruitment to lungs; 2) Our results were obtained with BcAU1054, a genomovar IIIB, PHDC lineage strain distinct from the BcJ2315 genomovar IIIA, ET12 lineage strain, suggesting that TecA is a virulence factor in all members of this species; 3) We demonstrate that TecA is a virulence factor in *Cftr*^F508del^ mice, which provide a useful CF animal model; 4) Our findings suggest that the presence of TecA in Bc is one reason why this species is most commonly responsible for cepacia syndrome(Loutet and Valvano 2010); 5) Our data support the idea that TecA is the founding member of a family of virulence factors in opportunistic bacterial pathogens, based on the finding that other opportunistic bacterial pathogens encode functional TecA orthologs (Aubert et al. 2016), including *Chryseobacterium indologenes* (Izaguirre-anariba and Sivapalan 2020) and *Ochrobactrum anthropi* (Hagiya et al. 2013). Finally, our results are significant because there are only a few examples of T6SS effectors that have demonstrated anti-host virulence factor activity in a mammalian infection model (Hachani, Wood, and Filloux 2016).

In an unexpected difference from what has been published with BcJ2315 (Xu et al.) we were not able to reproduce a requirement for pyrin in inflammatory cell recruitment to lungs during infection with either BcJ2315 or BcAU1054. Although BcJ2315 appears to be less virulent than BcAU1054, possibly due to the lack of O antigen (Aubert et al. 2016), this strain and BcAU1054 induced inflammatory cell recruitment to lungs independent of pyrin in our infection model. The TecA orthologs are 91% identical between the two strains (Aubert et al. 2016) and both BcJ2315 and BcAU1054 trigger assembly of pyrin inflammasomes in infected BMDMs. Other experimental differences could be responsible for the discrepancy. While we used o.p. infections with 5×10^7^ CFU, Xu et al. used intranasal infections with the 2-fold higher dose of 1×10^8^ CFU of BcJ2315 (Xu et al. 2014). Another difference is that the *Mefv^- /-^* mouse lines used in our study (Chae et al. 2011) and that of Xu et al (Xu et al. 2014) are both on the C57BL/6 background but independently generated and housed in different animal facilities. It is conceivable that mouse microbiota differences associated with the different animal facilities used in our study and that of Xu et al (Xu et al. 2014) are responsible for the discrepant results. It will be important to address this possibility in future studies.

Results of studies with BMDMs suggest that Bc infection can trigger T6SS-1-dependent activation of the NLRP3 (Rosales-Reyes et al. 2012; Valvano 2015) and non-canonical caspase-11 inflammasomes (Estfanous et al. 2021; Krause et al. 2018) in addition to the pyrin inflammasome. We did not detect significant amounts of IL-1β released from LPS-primed *Mefv*^-/-^ BMDMs infected with BcAU1054 or BcJ2315 (Fig. S3D, E), indicating that pyrin-independent inflammasomes are not activated during these in vitro infections. In a recent study, C57BL/6 *Gsdmd^-/-^* mice were infected o.p. with Bc strain K65-2 and the authors reported no difference in survival as compared to WT controls (Estfanous et al. 2021). These results are consistent with the idea that cleavage of GSDMD, which would occur downstream of NLRP3, caspase-11 or pyrin inflammasomes, is not essential for the virulence of Bc during lung infection as measured by a survival assay. However, this study reported that after BcK56-2 infection, *Gsdmd*^-/-^ mice had lower lung inflammation by H&E staining as compared to WT at 48 hours post challenge (Estfanous et al. 2021), suggesting that an inflammasome is contributing to immunopathology under these conditions. Additionally, GSDMD has a major role in restricting replication of BcK56-2 in macrophages, and at 48 hours post infection *Gsdmd*^-/-^ mice had higher BcK56-2 organ burdens as compared to WT (Estfanous et al. 2021). Understanding how different inflammasome pathways contribute to pathogenesis or host protection during Bc infections is an important goal in the field that will require additional research.

Although we demonstrated that pyrin is produced in BAL cells in response to BcAU1054 infection of WT mice, we have not attempted to determine if pyrin is activated by TecA in these cells. Aubert et al. obtained evidence that TecA triggers assembly of the pyrin inflammasome during systemic Bc infection in mice. In an intraperitoneal infection model, a Δ*tecA* mutant exhibited increased spleen burdens (day 4 post challenge) and lethality in WT mice as compared to the parental BcJ2315 strain (Aubert et al. 2016). However, in *Mefv*^-/-^ mice the Δ*tecA* mutant and parent exhibited similar lethality phenotypes, indicating that TecA triggers assembly of the pyrin inflammasome when infections are initiated systemically, although in this case it leads to a protective host response. It is possible that host phagocytes differentially control assembly of the inflammasome during Bc infections in distinct organs, such that this pathway is activated by pyrin in spleen but not lung.

How TecA promotes virulence during Bc lung infections remains to be determined. TecA does not increase BcAU1054 CFU in lungs suggesting that it does not directly counteract bactericidal activities of phagocytes when the infection is initiated by this route. T6SS-1 does not reduce phagocytosis of Bc although it can inhibit uptake of secondary targets 60 min after infection (Flannagan et al. 2012) which is consistent with the idea that the bacteria exploit the macrophage intracellular environment as an replicative niche. There is evidence that Bc can delay maturation of its phagosome in macrophages (Huynh et al. 2010; Lamothe et al. 2007) and recently this has been attributed to TecA (Walpole et al. 2020), but it remains unclear if this activity is essential for survival of the bacteria in host phagocytes. There is also evidence that T6SS-1 inhibits assembly of NADPH oxidase complexes on Bc-containing phagosomes in macrophages (Keith et al. 2009; Rosales-Reyes et al. 2012). However, the fact that only patients with defects in NADPH oxidase activity, such as CF or chronic granulomatous disease, are highly susceptible to Bc infections, strongly argues that these bacteria do not efficiently inhibit superoxide production in phagocytes. In a direct comparison of BcK56-2 and a *Burkholderia multivorans* clinical isolate in stimulating the oxidative burst in human macrophages, both strains induced the oxidative burst at roughly equivalent magnitudes (Assani et al. 2017). Thus, BcK56-2 which encodes *tecA,* does not appear to inhibit the oxidative burst more efficiently than a *B. multivorans* strain lacking this effector (Aubert et al. 2016). TecA also does not appear to increase dissemination of Bc from the lung, based on spleen or liver CFU of Bc, although this result may be hard to interpret if increased dissemination is balanced against decreased bacterial survival due to protective pyrin inflammasome responses in spleen or liver.

It is possible that TecA primarily promotes virulence during Bc infection by inducing lung immunopathology, as first observed by Aubert et al (Aubert et al. 2016). Immunopathology leading to lung failure could be the underlying basis for TecA-dependent weight loss and lethality in our infection model. A pyrin inflammasome-independent mechanism of immunopathology could be TecA-triggered cytokine or chemokine production, leading to increased numbers of F4/80^+^ inflammatory monocyte-derived macrophages in the Bc-infected lung, as we observed by IHC. CCL2 is produced by a variety of cells and signals for the recruitment of Ccr2^+^ inflammatory monocytes out of the bone marrow. We hypothesize that CCL2-elicited Ccr2^+^ inflammatory monocytes play a role in TecA-induced immunopathology as they may be the source of the F4/80^+^ cells in lung sections at 3 days post infection with BcAU1054 and these cells have been demonstrated to be detrimental to the host in other lung infection contexts (Heung and Hohl 2019). Studies using mice in which Ccr2^+^ inflammatory monocytes are reduced or can be depleted are underway to test this hypothesis.

Bacterial toxins that inactivate Rho GTPases can trigger activation of MAP kinases (MAPKs) in phagocytes (Warny et al. 2000), representing an alternative mechanism by which TecA could induce immunopathology. It has been shown that TcdA and TcdB trigger activation of MAPK pathways in monocytes and intestinal epithelial cells leading to release of IL-8 (Bobo et al. 2013; Warny et al. 2000). The activation of MAPK MK2 (pMK2) has been demonstrated in epithelial cells as well as in mouse and hamster intestines (Bobo et al. 2013). IL-8 is a potent neutrophil chemoattractant and secretion of IL-8 in the gut mucosa leads to injury (Bobo et al. 2013; Warny et al. 2000). Interestingly, IL-8 has also been shown to be secreted at higher levels in CF immune cells as compared to non-CF and is present at higher levels in CF patient BAL samples suggesting that there are alterations in MAPK signaling in CF (Nakamura et al. 1992; Zaman et al. 2004).

Bc infection in *Cftr*^F508del^ mice does not recapitulate all aspects of the disease in CF patients (e.g., lack of preceding chronic infection and lung necrosis in the context of cepacia syndrome), but we consider it a reasonable and useful CF animal infection model. This model can be used to uncover the virulence mechanisms of TecA. In addition, this model can be used to better understand how immune deficiencies in CF contribute to susceptibility to Bc lung infection. For example, does dysbiosis of the microbiota in CF contribute to risk for Bc lung infection? Finally, this model can be used to understand mechanisms of protective immunity to Bc, resulting in insights toward development of vaccines and immunotherapeutics to treat or prevent Bc infections in the CF patient population.

## Materials & Methods

### Bacterial strains

A list of all *Burkholderia cenocepacia* (Bc) strains used in this study can be found in Table 1. Bc overnights were grown in Luria-Bertani (LB) media shaking cultures at 37°C. The deletion mutants BcAU1054 Δ*tecA* and Δ*hcp* were made by allelic exchange (Perault et al. 2020). For each mutant, ∼500 base pairs 5’ to and including the first three codons of the gene to delete were fused to ∼500 base pairs 3’ to and including the last three codons of the gene by splicing by overhang extension PCR. Fusion products were cloned into the allelic exchange vector pEXKm5 (López et al. 2009) and plasmids were conjugated into BcAU1054 using *E. coli* strain RHO3. Merodiploids were selected on 250 µg/mL kanamycin and grown for 4 h at 37°C with aeration in YT broth (10 g/L yeast extract, 10 g/L tryptone), subcultured 1:1000 fresh YT broth, and grown overnight at 37°C with aeration. Following overnight growth, cells that had resolved the integrated plasmid were selected on YT agar (1.5% agar) containing 25% sucrose and 100 µg/mL 5-bromo-4-chloro-3-indoxyl-β-D-glucuronide and incubated at 30°C, and deletions were confirmed via PCR and sequencing of the regions spanning the deletions. Complementation of the cloned wild-type (WT) gene or *tecA_C41A_,* genes were done at transposon insertion sites in the Bc chromosome (*att*Tn7 sites) using the pUC18T-mini-Tn7T suite of plasmids (Choi et al. 2005) and were selected by kanamycin resistance. Complementation sequences were cloned into pUC18T-mini-Tn7T-Km containing the constitutive ribosomal S12 subunit gene promoter of *Burkholderia thailandensis* E264 immediately 5’ to the multiple cloning site (plasmid pUCS12Km, (Anderson, Garcia, and Cotter 2012)). The *tecA_C41A_* sequence was generated using the Agilent QuikChange II Site-Directed Mutagenesis Kit (#200523). Complementation cassettes were delivered to the BcAU1054 *att*Tn7 site via triparental mating with *E. coli* RHO3 strains harboring pUCS12Km-*tecA_WT_*/pUCS12Km-*tecA_C41A_* and the transposase helper plasmid pTNS3, and BcAU1054 exconjugants containing these cassettes were selected on agar containing 250 µg/mL kanamycin. All Bc strains used in this study are ampicillin resistant.

### Ethics statement

Studies requiring mice for isolation of bone barrow and live mice for infections were carried out in accordance with a protocol that adhered to the Guide for the Care and Use of Laboratory Animals of the National Institutes of Health (NIH) and was reviewed and approved (approval #00002184) by the Institutional Animal Care and Use Committee at Dartmouth College. The Dartmouth College animal program is registered with the U.S. Department of Agriculture (USDA) through certificate number 12-R-0001, operates in accordance with Animal Welfare Assurance (NIH/PHS) under assurance number D16-00166 (A3259-01) and is accredited with the Association for Assessment and Accreditation of Laboratory Animal Care International (AAALAC, accreditation number 398). Age-matched, sex-matched, and/or littermate controls were used when appropriate.

### Mouse strains

Wild-type (WT) C57BL/6J (stock# 000664) were purchased from Jackson Laboratories at 7 weeks of age and rested for a week in our mouse facility before infection. Pyrin-knockout mice (*Mefv^-/-^*) on the C57BL/6 background were obtained from Jae Chae and Daniel Kastner at the NIH (Chae et al. 2011) and bred in the mouse facilities at Dartmouth. Mice with the *Cftr^F508del^* mutation on the C57BL/6 background were obtained from Case Western Reserve University’s Cystic Fibrosis Mouse Models Core and bred at Dartmouth. *Mefv^-/-^* and *Cftr^F508del^* mice were crossed to obtain breeding pairs that were *Cftr^F508del^ Mefv^+/-^* and *Cftr^F508del^ Mefv^-/-^*. The resulting pups were genotyped and used for infections.

### Cell culture

Bone marrow derived macrophages (BMDMs) were cultured from bone marrow of mice and cultured as described previously(Brodsky et al. 2010; Chung et al. 2016). After 7 days of differentiation the BMDMs were seeded at a density of 0.8×10^6^ cells/well in 6-well plates in MGM 10/10 media containing Dulbecco’s Modified Eagle Medium (DMEM) + Glutamax (Gibco^®^) containing 10% Fetal Bovine Serum (FBS) (GE^®^), 10% L929 cell-conditioned media, 1 mM sodium pyruvate (Gibco^®^), 10 mM HEPES (Gibco^®^) and divided into 6-well plates at a density of 0.8 x10^6^ cells/well in a total volume of 3 mL. The BMDMs were primed with 100 ng/mL O26:B6 *Escherichia coli* LPS (Sigma^®^) and incubated overnight at 37°C with 5% CO_2_.

### Macrophage infections

Overnight (16-hour) cultures of *B. cenocepacia* were subcultured 1:100 in fresh LB on infection day and shaken at 37°C until the cultures reached mid-log phase (OD_600_ = 0.300). Cultures were then pelleted, the LB was removed and the bacteria were resuspended in PBS to the original volume. The bacterial suspensions were then diluted to an MOI of 20 in warmed, serum free MGM 10/10. As a positive control for pyrin inflammasome activation, the glucosyltransferase toxin from *Clostridium difficile* TcdB (List Biological Laboratories, Inc.) was also diluted to 0.1μg/ml in warmed serum free MGM 10/10. The BMDMs were washed once in warm 1x PBS and 3ml fresh serum free MGM 10/10 was added with bacteria and TcdB to the appropriate wells. The plates were then centrifuged for 5 minutes at 1,000rpm to bring the bacteria and the cells in contact on the bottom of the wells. The plates were incubated at 37°C with 5% CO_2_ for 90 minutes. Cell supernatants were collected for cytokine ELISAs and lactate dehydrogenase (LDH) assays. The BMDMs were lysed using mammalian protein extraction reagent (M-PER, Thermo Scientific^®^) with added cOmplete, Mini (Roche^®^) protease inhibitor and PhosSTOP (Roche^®^) phosphatase inhibitor.

### Protein analysis by SDS-PAGE and Western blot

5 - 10μg of protein from the cell lysates were run on 4-12% NuPAGE Bis-Tris SDS-PAGE gels (Invitrogen by ThermoFisher Scientific^®^) and transferred to PVDF membranes (ThermoFisher Scientific^®^) using an iBlot 2 Gel Transfer Device (life technologies). Membranes were blocked in 5% non-fat dairy milk and incubated with primary antibodies overnight. The primary antibodies used are as follows; rabbit-anti-mouse MAb total pyrin antibody (abcam^®^, ab195975), rabbit-anti-mouse MAb phospho-Serine 205 (abcam^®^, ab201784), rabbit-anti-mouse/human IL-1β (Cell Signaling^®^, #12242), rabbit-anti-mouse/human polyclonal β-actin (Cell Signaling^®^, #4967). HRP-conjugated anti-rabbit antibody (Jackson laboratory) was used as a secondary. Proteins were visualized using chemiluminescent detection reagent (GE Healthcare^®^) on an iBright FL1500 (ThermoFisher Scientific^®^).

### IL-1β quantification

IL-1β in BMDM supernatants was quantified using a murine ELISA kit (R&D Systems^®^, MLB00C) following the manufacturer’s instructions.

### Tail genotyping

Genomic DNA was isolated from mouse tail pieces (clipped at time of weaning) by digestion overnight at 55°C in 500μl lysis buffer (100mM Tris-HCl, pH 8.5, 5mM EDTA, 200mM NaCl, 0.2% SDS, 100μg/ml Proteinase K (VWR)). The supernatant was collected by centrifugation (14,000rpm, 10 minutes, room temperature) and DNA was precipitated with 500μl 100% isopropanol. After centrifugation (14000rpm, 10 minutes, room temperature), the pellets were washed once with 70% ethanol, air-dried for 20 minutes, and dissolved in 200μl TE buffer at 55°C for 20 minutes. PCR was performed with the following primers: CTGCCCAGAGAAAGGTGATT WT F (ATCAAAGAAAATATCATCTTT), *Cftr* WT R (GGACGGTATCATCCCTGAAA), *Cftr* F508del F (ATCAAAGAAAATATCATTGGT), *Cftr* F508del R (ATGGACGGTATCATCCCTGA), Internal control F (CTAGGCCACAGAATTGAAAGATCT), Internal control R (GTAGGTGGAAATTCTAGCATCATCC), *Mefv* WT F (TGGAAATGGGAGTCCAGAAA), *Mefv* WT R (ACCTACCTGTGGGGTCACTG), *Mefv* KO F (GGGGGAACTTCCTGACTAGG), and *Mefv* KO R (CTGCCCAGAGAAAGGTGATT). To genotype the mouse *Cftr* gene each tail gDNA sample was amplified by PCR in two reactions: one with WT Cftr primers + internal control and another with F508del Cftr primers + internal control. To genotype the murine *Mefv* gene each tail gDNA sample was amplified by PCR in two reactions: one with WT Mefv primers and another with the Mefv KO primers. PCR products were analyzed by agarose gel electrophoresis.

### Mouse infections

Overnight (16-hour) cultures of *B. cenocepacia* were subcultured 1:100 in fresh LB on infection day and shaken at 37°C until the cultures reached mid-log phase (OD_600_ = 0.300). To prepare the cultures for inoculation the appropriate volume of culture was centrifuged at 14,000 rpm for 5 minutes. The LB was removed, the pellets were combined using a small volume of sterile 1x PBS if the original culture volume exceeded the volume of one tube and the cultures were pelleted again at 14,000 rpm for 5 minutes. The remaining supernatant was carefully aspirated and the pellet was resuspended in sterile 1x PBS to final volume for 50μl/dose. The inoculum was serially diluted, plated on LB plates, grown overnight at 37°C and counted to ensure the dose was correct for each Bc strain used.

Both male and female mice 6-12 weeks of age were anesthetized with isoflurane and inoculated via the o.p. route with one 50μl dose or PBS (mock) of 5×10^7^ CFU of Bc using a pipette. Mice were weighed immediately following instillation and weight was monitored at 24-hour intervals from time of instillation throughout the experiment. Mice were monitored for survival for 14 days and were checked 3 times a day and euthanized if moribund. At indicated time points after infection mice were euthanized with CO_2_ or Euthasol with the help of a veterinarian. Tissues including lungs, bronchoalveolar lavage fluid (BALF), liver and spleen were collected. BALF was collected by inserting a needle into the trachea, removing the lungs and trachea together and filling the lungs with sterile 1x PBS once with 1ml and twice after with 500μl, slowly aspirating and massaging the lungs to recover as much BALF as possible. The cells in the BALF were counted and aliquots of each sample were lysed with either NP-40 (50mM Tris, pH 8.0, 150mM NaCl, 1% NP-40) to release intracellular bacteria for subsequent serial dilutions and plating for CFU, or M-PER (with protease and phosphatase inhibitors as described above) to generate whole cell lysates. The protein concentration of the BALF lysates were quantified, ran on SDS-PAGE gels, and used for Western blotting.

To prepare Bc antisera, a C57BL/6J mouse was infected as above except the route was intranasal and the dose was 1×10^8^ CFU of BcAU1054. At day 15 the infection was repeated with a dose of 5×10^7^ CFU of BcAU1054. At day 28 the mouse was euthanized and serum was collected and used for IHC along with control serum collected from a naïve C57BL/6J mouse.

### Lung fixation, paraffin embedding, H&E and IHC staining

Lungs were harvested and the left lungs were used for organ burden (see below) and right lungs were inflated with 10% neutral buffered formalin (Fisher Scientific) and left submerged in formalin for at least 24 hours. The right lungs were then dissected into the four different lobes and placed in cassettes submerged in 70% ethanol. The four right lobes for each lung were paraffin embedded and sectioned by the DHMC pathology core. The four lobe sections for each right lung arranged side-by-side on a slide were then stained with hematoxylin and eosin (H&E) for histopathology or were stained to detect specific cell subsets with one of the following antibodies suitable for IHC: Ly6G (abcam^®^ ab238132), CD11b (abcam^®^ ab133357), F4/80 (D2S9R) (Cell Signaling^®^ #70076), followed by diaminobenzidine (DAB)-conjugated secondary antibody. Alternatively, sections were stained with BcAU1054 antisera and DAB-conjugated secondary antibody to detect the bacteria.

### Microscopy & pathological analysis

H&E and IHC images of the four lobe sections for each right lung were captured on a Zeiss Axioskop 2 light microscope with a SPOT Insight sCMOS camera (SPOT Advanced Software) maintained by the Dartmouth microscopy core at 5X, 10X, 20X and 40X magnification (Plan Neofluar objectives). H&E images of the four lobes from each right lung were blinded and analyzed by Joseph Schwartzman using standard pathological criteria for inflammation and damage.

Semi-quantitative H&E scoring and quantitative IHC staining image analysis were performed by HistoWiz. Slides were scanned and the blinded H&E images of the four lobes from each right lung were subjected to semi-quantitative scoring of inflammation and damage as described (Fukushi et al. 2011) Lesion severity was evaluated on a minimal, mild, moderate, and marked scale and lesion distribution was classified as focal, multifocal, or diffuse (Table S1). Images of the four lung lobe sections from each right lung on a scanned IHC slide were subjected to analysis and cellular quantification was performed to identify total cells and percentages of DAB positive cells classified by weak, medium, or intense stain. Values were normalized by H-Score calculated as 1 x (% weak stain) + 2 x (% medium stain) + 3 x (% intense stain).

### Organ burden

Organs were collected in stomacher bags, weighed, and placed on ice. 5ml of sterile 1x PBS was added to full (left and right) lungs and livers and 2.5ml of sterile 1x PBS was added to spleens and left lungs. Organs were homogenized by rolling out by hand and then by use of a Stomacher^®^ 80 Biomaster (Seward) on high setting for 2 minutes. For lungs this homogenization method was done twice to increase homogenization of the tissue. The homogenates were then serially diluted in sterile, 1x PBS and plated on LB or LB plates with 100μg/ml ampicillin. The plates were incubated at 37°C and the colonies were counted at least 24 hours later. Data are displayed as CFU per gram of tissue.

### Statistical analysis

GraphPad Prism was used to perform statistical analyses. Cytokine assays were analyzed by the Mann-Whitney test or one-way ANOVA (Tukey’s multiple comparisons test). Error bars shown represent standard deviation. Analysis of the weight loss data were performed by Thomas H. Hampton in the R statistical programming language including ggplot2. Asymptotic Wilcoxon-Mann-Whitney tests were performed using the coin package. Weight loss data was also analyzed using Welch’s T test. Additional details for each experiment can be found in the figure legends. Survival data were analyzed using the Log-Rank (Mantel-Cox) test.

---

**Fig. S1.**
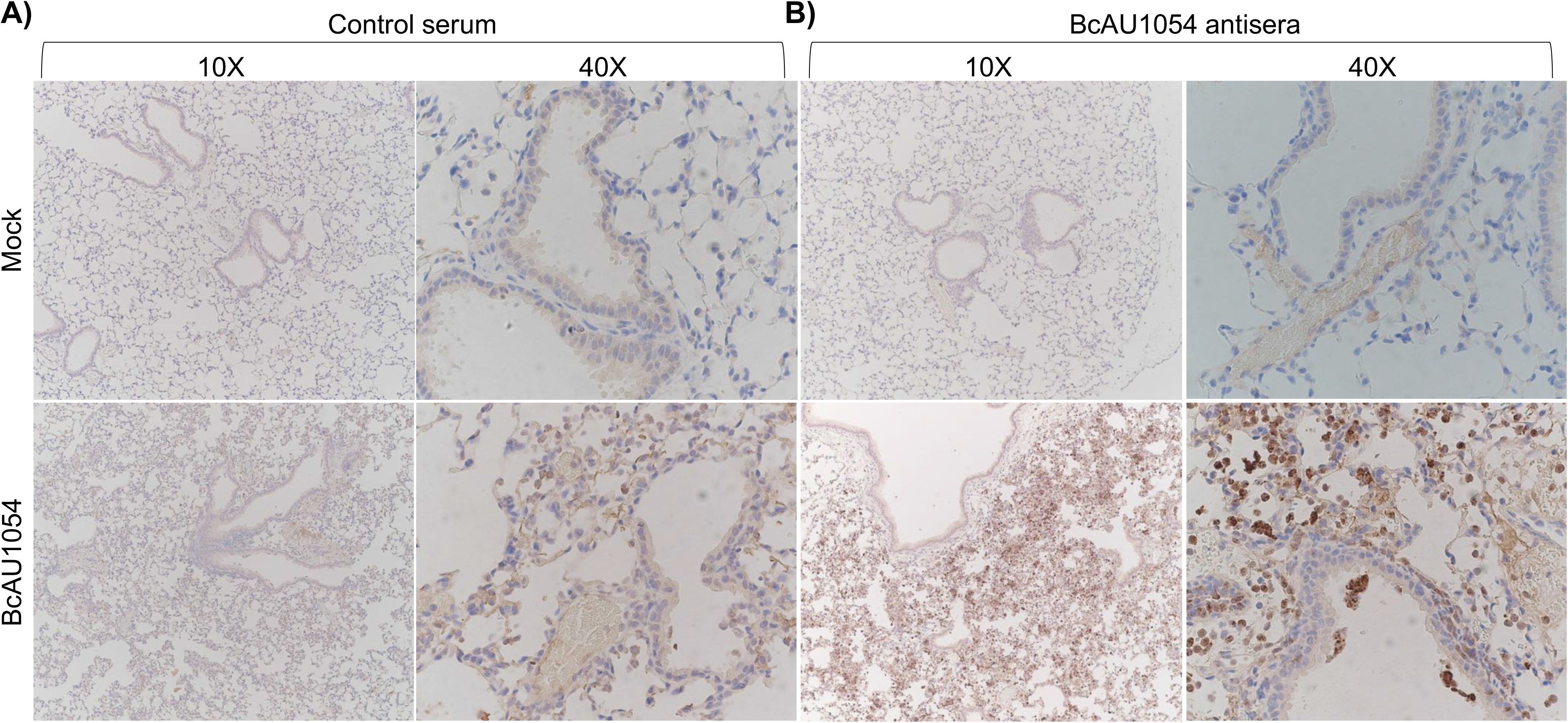
IHC staining of WT mouse lung sections infected with BcAU1054. IHC was performed with A) control serum or B) BcAU1054 antisera on fixed right lung lobe sections from a WT mouse left uninfected (mock) or infected by o.p. instillation with 5×10^7^ CFU of BcAU1054 for 12 hours. Representative images captured by light microscopy are shown at 10X or 40X magnification.

**Fig. S2.**
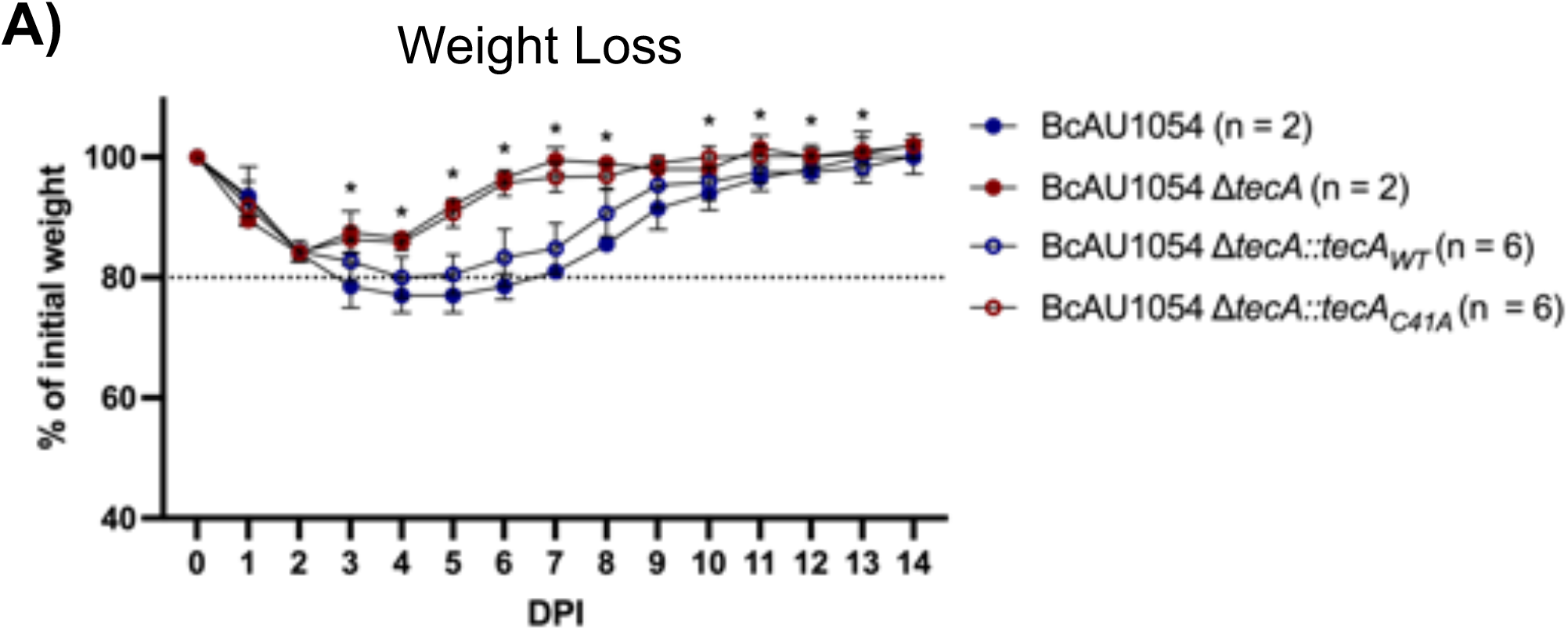
Enzymatic activity of TecA is necessary for exacerbated weight loss in WT mice. WT mice were infected o.p. with 5×10^7^ CFU of BcAU1054, Δ*tecA*, Δ*tecA::tecA_WT_*, or Δ*tecA::tecA_C41A_* and weighed on D_0_ and daily for 14 DPI. Data are represented as percent of initial weight. Days on which mice infected with Δ*tecA::tecA_WT_* or Δ*tecA::tecA_C41A_* differed significantly (n=6, p < .05, Welch’s T test) are marked with an asterisk. Results are from one experiment in which none of the mice died, n = total mice infected.

**Table S1.** Summary of semi-quantitative scoring of lung inflammation and damage.

| Summary of semi-quantitative scoring of lung inflammation and damage. |  |  |  |
| --- | --- | --- | --- |
| Timepoint | Condition | Bronchioles | Alveoli |
| 12hpi | Mock | Normal | Normal |
|  | BcAU1054 | Intraluminal neutrophilic and mononuclear cell infiltrate, mild, multifocal | Neutrophilic and histiocytic inflammatory infiltrate, multifocal, mild<br>Alveolar collapse: mild, multifocal<br>Alveolar septal thickening: mild, multifocal |
| | Bc AU1054 ΔtecA | Intraluminal neutrophilic and mononuclear cell infiltrate, moderate, multifocal | Neutrophilic and histiocytic inflammatory infiltrate, moderate, multifocal<br>Alveolar collapse: mild, multifocal<br>Alveolar septal thickening: mild, multifocal |
| 3dpi | Mock | Normal | Normal |
|  | BcAU1054 | Intraluminal neutrophilic and mononuclear cell infiltrate, mild, multifocal | Neutrophilic and histiocytic inflammatory infiltrate, marked, multifocal<br>Alveolar collapse: moderate, multifocal<br>Alveolar septal thickening: moderate, multifocal |
| | Bc AU1054 ΔtecA | Intraluminal neutrophilic and mononuclear cell infiltrate, mild, multifocal | Neutrophilic and histiocytic inflammatory infiltrate, marked, multifocal<br>Alveolar collapse: marked, multifocal<br>Alveolar septal thickening: marked, multifocal |

**Fig. S3.**
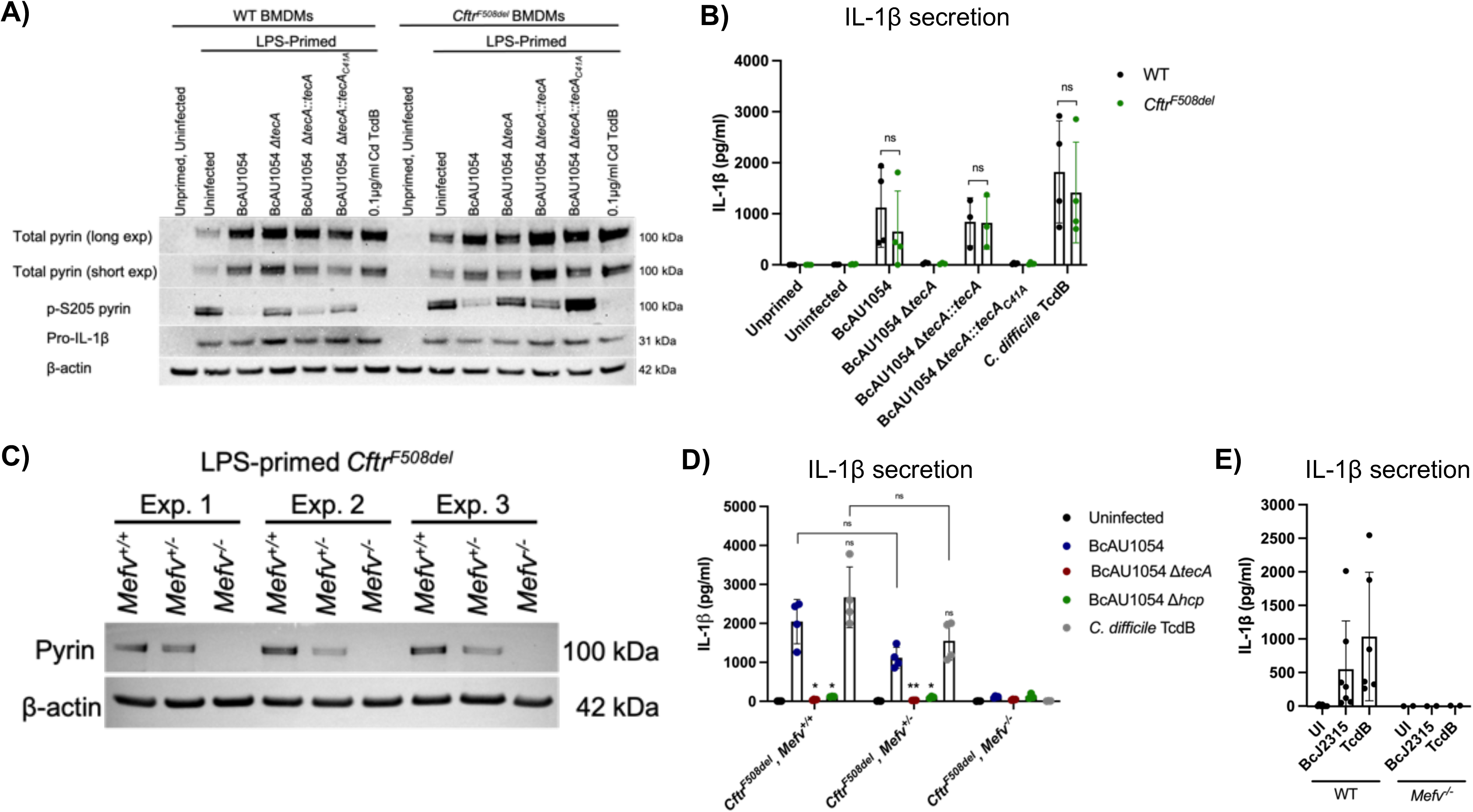
Bc TecA activates the pyrin inflammasome in WT and *Cftr^F508del^* BMDMs. A) Whole cell lysates from WT and *Cftr^F508del^* BMDMs left unprimed or primed with LPS and infected at MOI 20 with BcAU1054, Δ*tecA*, Δ*tecA::tecA_WT_*, Δ*tecA::tecA_C41A_* or intoxicated with TcdB for 90 minutes were analyzed by western blotting with antibodies to total pyrin, p-S205 pyrin, pro-IL-1β or β-actin (loading control). B) IL-1β quantified by ELISA in cell supernatant collected from WT and *Cftr^F508del^* BMDMs left unprimed or primed with LPS and infected with Bc or intoxicated with TcdB as in A). Data shown is from at least three independent experiments. Error bars represent standard deviation. Significance determined by two-way ANOVA. C) Whole cell lysates from *Cftr^F508del^ Mefv^+/+^*, *Cftr^F508del^ Mefv^+/-^*, and *Cftr^F508del^ Mefv^-/-^* BMDMs primed with LPS were analyzed by western blotting with antibodies to total pyrin and β-actin (loading control). D) IL-1β quantified by ELISA in cell supernatant collected from *Cftr^F508del^ Mefv^+/+^*, *Cftr^F508del^ Mefv^+/-^*, and *Cftr^F508del^ Mefv^-/-^* BMDMs primed with LPS and infected with BcAU1054, Δ*tecA* or Δ*hcp* or intoxicated with TcdB for 90 minutes. Data shown is from at least three independent experiments. Error bars represent standard deviation. Significance in D) determined by one-way ANOVA (Tukey’s multiple comparisons test) between conditions within cell genotypes and between cell genotypes by Mann-Whitney test. E) IL-1β quantified by ELISA in cell supernatant collected from WT and *Mefv^-/-^* BMDMs primed with LPS and infected with BcJ2315 or intoxicated with TcdB for 90 minutes. Data shown is from at least three independent experiments. Error bars represent standard deviation.

**Fig. S4.**
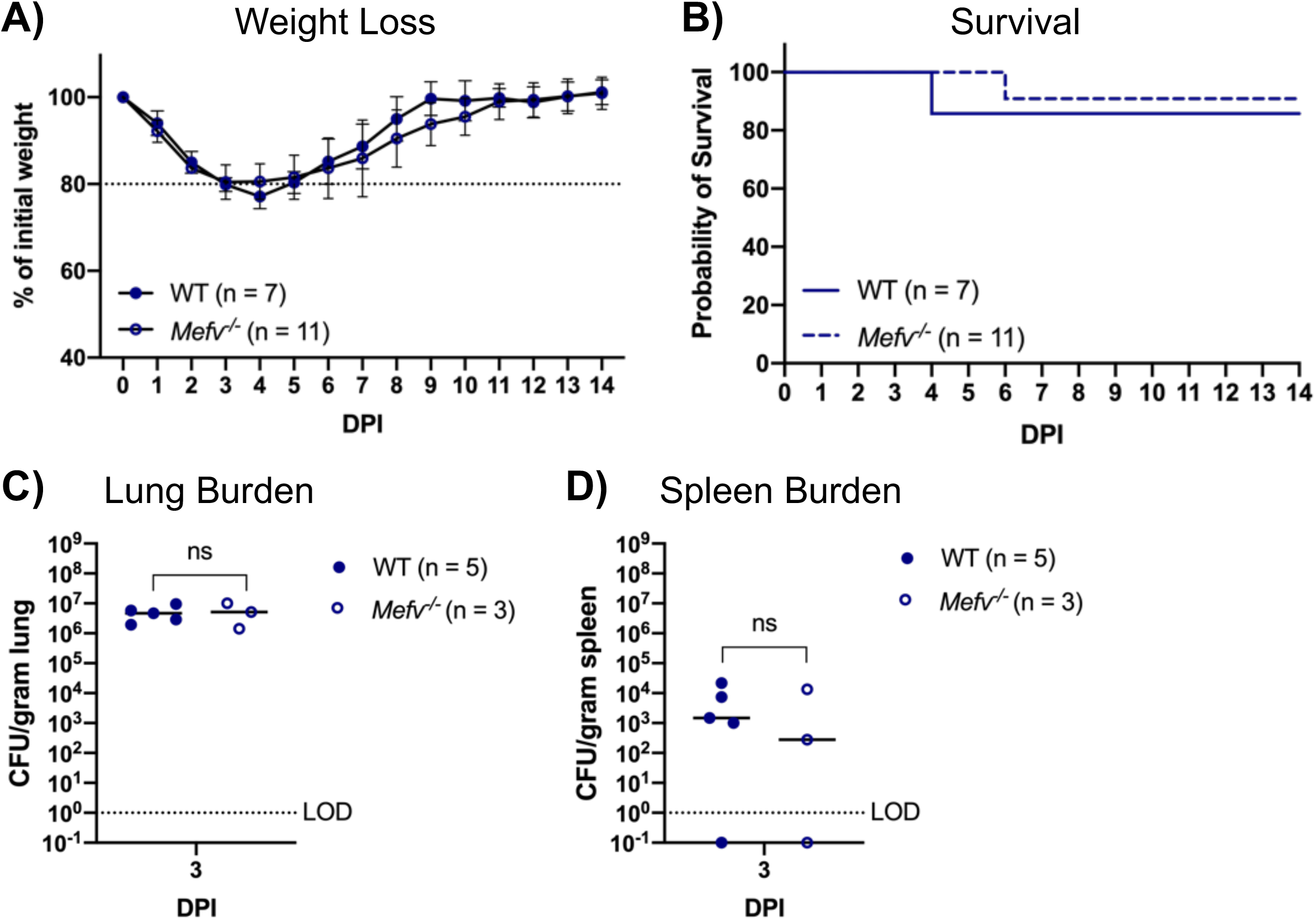
Pyrin is dispensable for BcAU1054 virulence in WT mice. A) Weight loss of WT and *Mefv^-/-^* mice infected o.p. with 5×10^7^ CFU of BcAU1054. Mice were weighed on D_0_ and daily for 14 DPI. Results were pooled from 2 independent experiments, n = total mice infected. Data are represented as percent of initial weight. B) Percent survival monitored for 14 DPI of WT and *Mefv^-/-^* mice infected with BcAU1054. Results were pooled from 2 independent experiments, n = total mice infected. ns, not significant by Log-Rank (Mantel-Cox) test. C, D) Lung and spleen burden at the indicated day post infection (DPI) in WT and *Mefv^-/-^* mice infected with BcAU1054. Results from one experiment, each point represents a value obtained from an individual mouse. Data are represented as CFU per gram of tissue. The mice used in these experiments were not controlled using littermates or co-housing.

**Figure S5.**
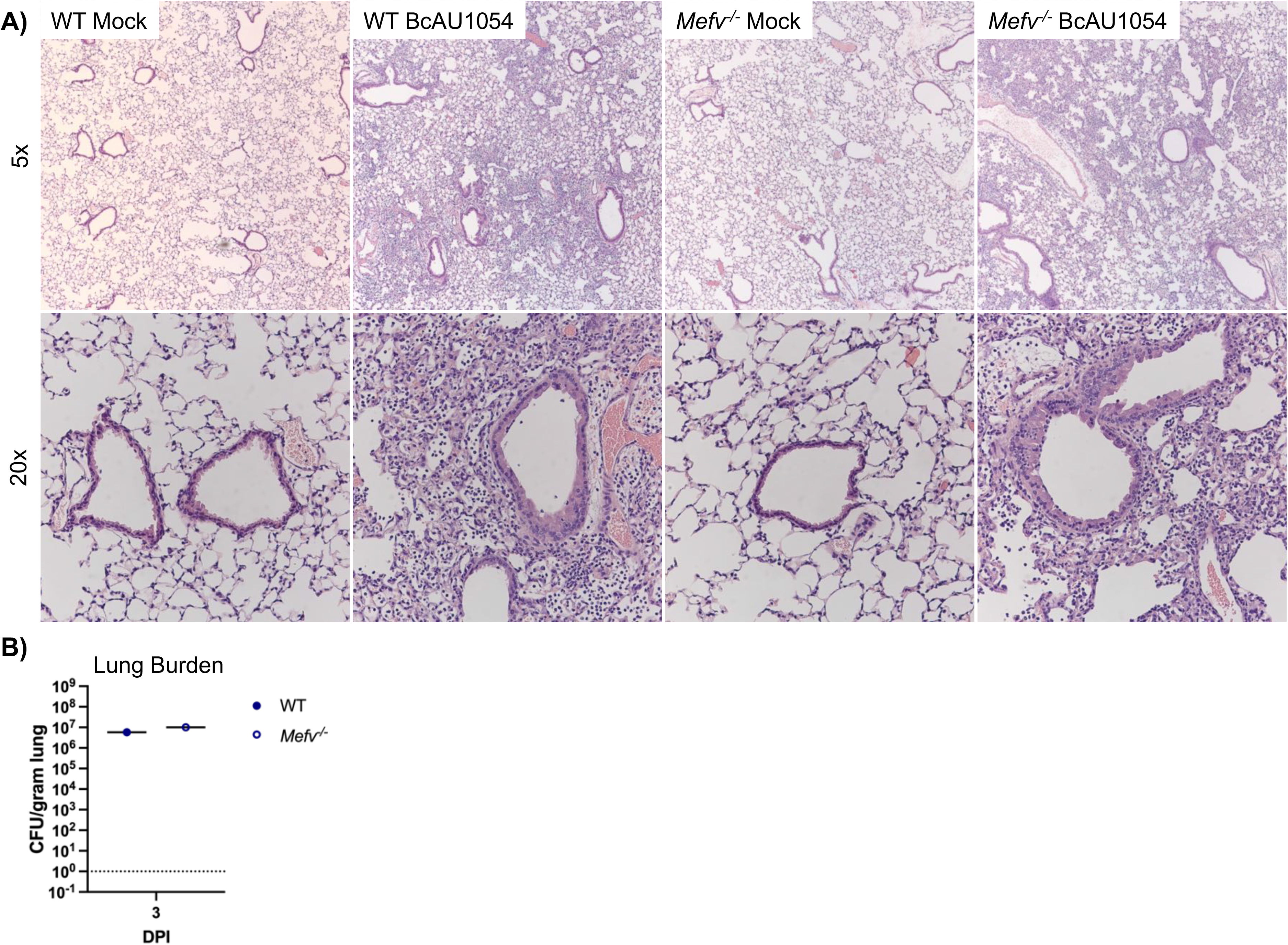
Pyrin is dispensable for lung inflammation in WT mice infected with BcAU1054. A) Representative H&E images of the four right lung lobe sections from a WT or *Mefv^-/-^* mouse uninfected (mock) or infected o.p. with 5×10^7^ CFU BcAU1054 at 3dpi. Light microscopy images are shown at 5X and 20X magnification. B) Lung burdens of BcAU1054 in infected left lungs of mice analyzed in A). Data are represented as CFU per gram of lung.

**Figure S6.**
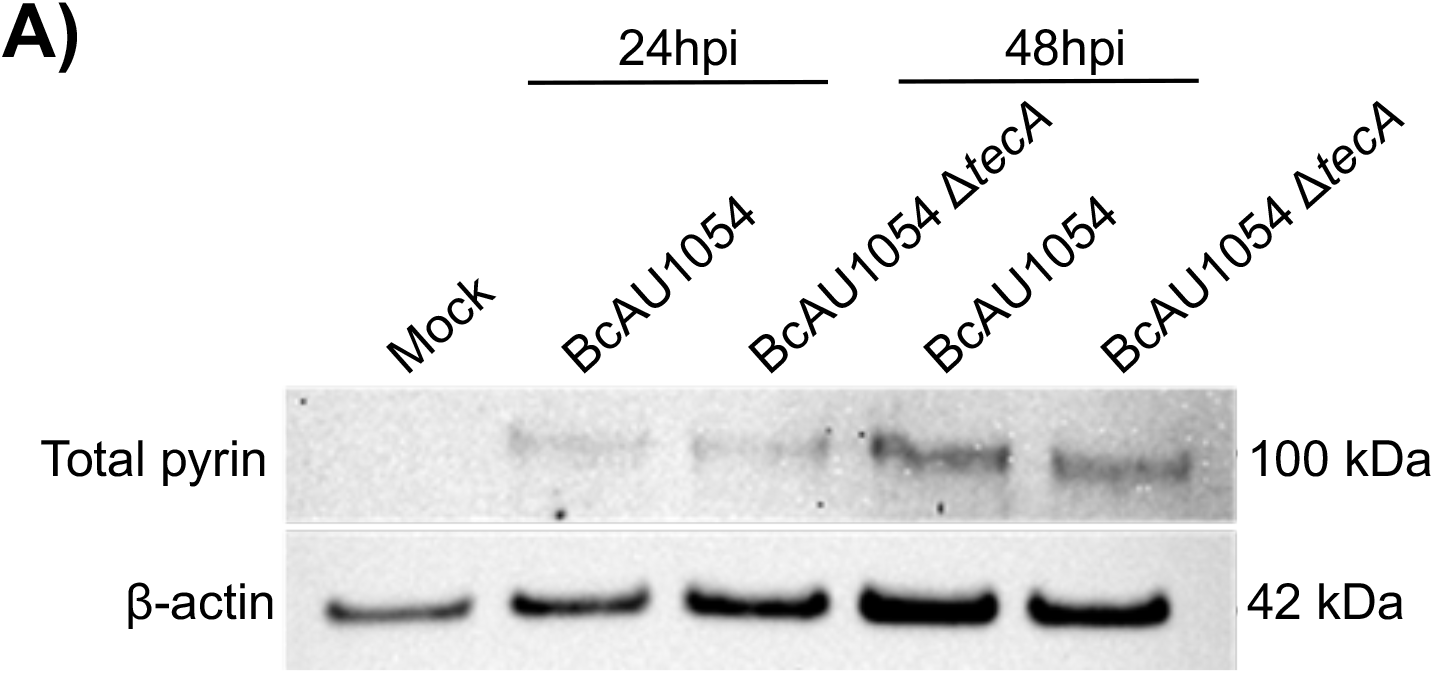
Pyrin protein levels in BAL cell lysates from infected WT mice. WT mice were left uninfected (mock) or infected o.p. with 5×10^7^ CFU of BcAU1054 or Δ*tecA* and BALF was collected at indicated time points. Lysates from BAL cells were normalized for total protein and immunoblotted for total pyrin or β-actin (loading control).

**Figure S7.**
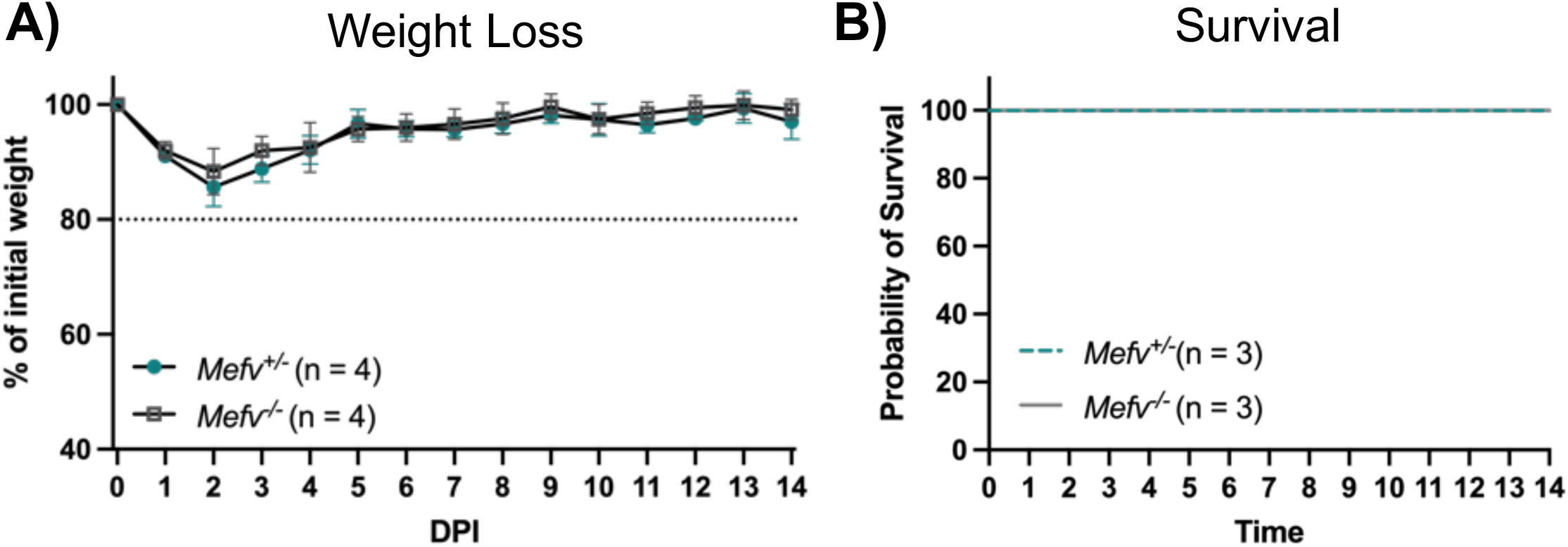
Pyrin is dispensable for weight loss in WT mice infected with BcJ2315. A) Weight loss of *Mefv^+/-^* and *Mefv^-/-^* mice infected o.p. with 5×10^7^ CFU BcJ2315. Mice were weighed on D_0_ and daily for 14 DPI. Data are represented as percent of initial weight. B) Percent survival monitored for 14 DPI of *Mefv^+/-^* and *Mefv^-/-^* mice infected with BcJ2315. Results shown are from one experiment, n = total mice infected for both A) and B).

**Figure S8.**
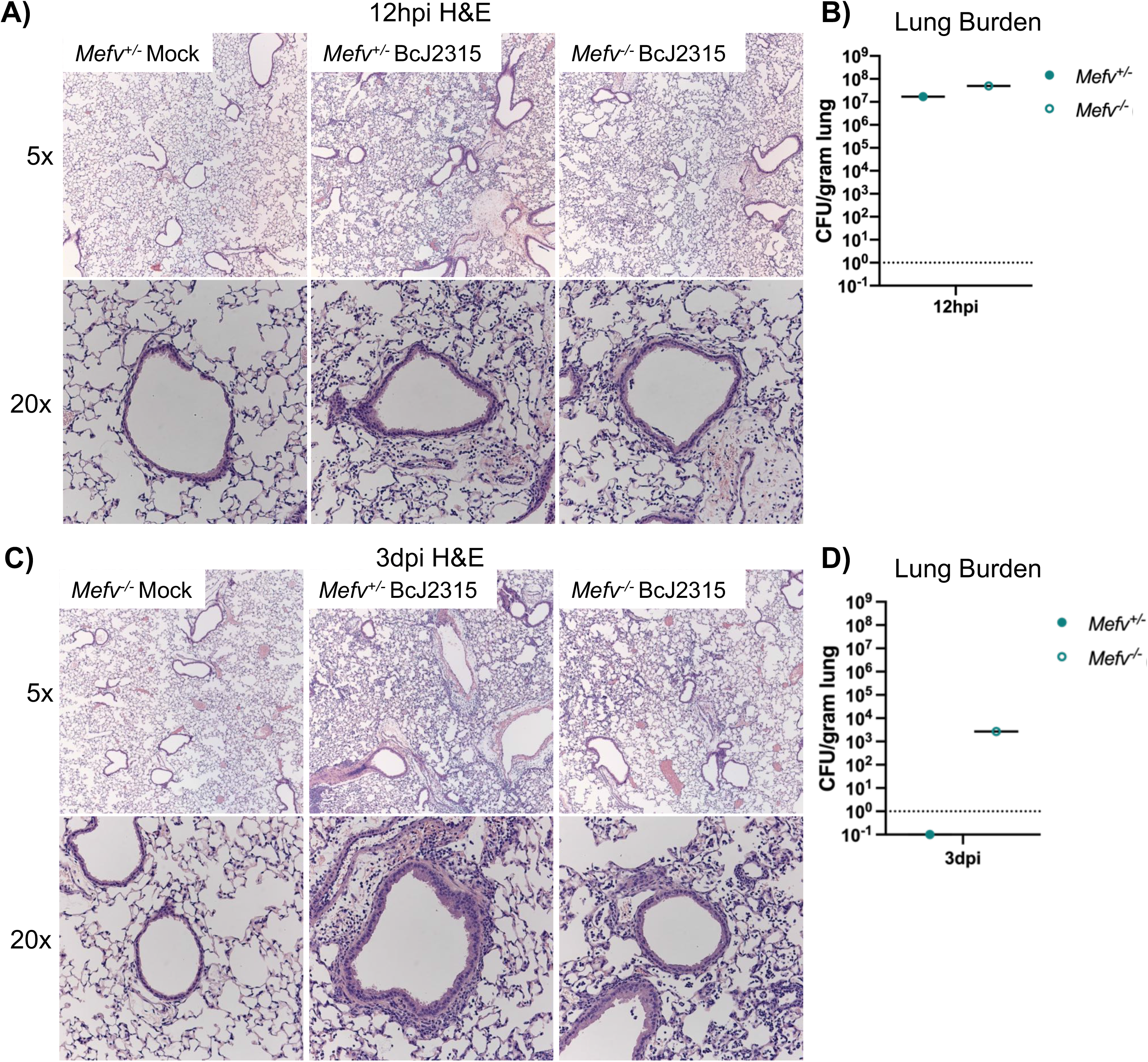
Pyrin is dispensable for lung inflammation in WT mice infected with BcJ2315. Representative H&E images of the four right lung lobe sections from a *Mefv^+/-^* or *Mefv^-/-^* mouse left uninfected (mock) or infected with 5×10^7^ CFU BcJ2315 for A) 12 hours or C) 3 days. Light microscopy images are shown at 5X and 20X magnification. Burdens of BcJ2315 in the infected left lungs analyzed in A) and C) are shown in B) and D) respectively. Data are represented as CFU per gram of lung.

## Author contributions

N.A.L. did all experiments and contributed to writing this manuscript. A.I.P. and P.A.C. provided *Burkholderia cenocepacia* AU1054 strains and edited this manuscript. C.A. H. provided *Cftr*^F508del^ mice and edited this manuscript. J.D.S. is a clinical microbiologist and pathologist who reviewed the H&E slides. T.H.H. did statistical analysis for the mouse weight loss data. J.B.B. provided funding, was integral in experimental design and helped write and edit this manuscript.

## Acknowledgements

Jane Jones and Robb Cramer provided instruction and advice on o.p. infections. Ann Lavanway was instrumental in helping with microscopy. *Burkholderia cenocepacia* J2315 was generously provided by George O’Toole, Jr. The Pathology Core staff especially, Scott Palisoul and Rebecca O’Meara, helped with H&E and IHC. We also thank Wei Wang and the CCMR staff especially Kevin Legace, Hannah Salamy and Eric DuFour for assistance with mouse husbandry and experimentation.

